# A rhythmically pulsing leaf-spring nanoengine that drives a passive follower

**DOI:** 10.1101/2021.12.22.473833

**Authors:** Mathias Centola, Erik Poppleton, Martin Centola, Julián Valero, Petr Šulc, Michael Famulok

## Abstract

Molecular engineering seeks to create functional entities for the modular use in the bottom-up design of nanoassemblies that can perform complex tasks. Such systems require fuel-consuming nanomotors that can actively drive downstream passive followers. Most molecular motors are driven by Brownian motion, but the generated forces are scattered and insufficient for efficient transfer to passive second-tier components, which is why nanoscale driver-follower systems have not been realized. Here, we describe bottom-up construction of a DNA-nanomachine that engages in an active, autonomous and rhythmical pulsing motion of two rigid DNA-origami arms, driven by chemical energy. We show the straightforward coupling of the active nanomachine to a passive follower unit, to which it then transmits its own motion, thus constituting a genuine driver-follower pair. Our work introduces a versatile fuel-consuming nanomachine that can be coupled with passive modules in nanoassemblies, the function of which depends on downstream sequences of motion.

## Introduction

Active mechanical motion of nanoscale objects is paramount for the bottom-up construction of bio- or technomimetic nanomechanical machines^1–6^ that can perform tasks like pumping^7^, walking^8^, transduction or sensing of molecules or signals^9^, or any process involving motion^10–11^. Both in the nano- and in the macroscopic world these processes require fuel-powered engines that perform periodically repeating rhythmic motion. Impressive examples of synthetic pumping, rotating, or moving fuel-driven nano-devices exist^12–16^ but the creation of engines that generate active rhythmic motion at the nanoscale, driven by chemical fuel, remains challenging^17–19^. Here we report a bio-hybrid nanoengine (NE) that rhythmically pulses by consuming nucleoside triphosphates as fuel to build up potential energy stored as spring-tension in a compliant flexure mechanism, followed by active relaxation. The device consists of two rigid DNA origami-arms connected in a V-shape by a bendable flexure at the pointed end, and a double-stranded DNA (dsDNA) sequence that spans the diverging ends^20^. One origami-arm has a T7-RNA polymerase (T7RNAP) covalently attached near a T7 promotor sequence in the dsDNA, allowing its use as a transcription template to create traction^21–23^. Upon transcription, the stationary T7RNAP pulls the dsDNA like a rope and drags the opposing origami-arms towards itself, which builds up spring tension at the compliant flexure that is released when the polymerase reaches a termination sequence near the opposite origami arm allowing the structure to return to its equilibrium position. This design leads to a continuous rhythmic flapping motion the frequency of which can be tuned by slight changes in the transcribed dsDNA sequence. Coarse grained molecular dynamics simulations illustrate the mechanical properties of various nanoengine designs and confirm our experimental results. In a prototypical application we show that the engine acts as a mechanical driver that can be coupled to a passive follower unit to which it then can actively transfer its motion, opening ground-breaking opportunities for its future use to drive more complex nanomachines, similar to the balance wheel in a watch or in Leonardo da Vinci’s self-propelled cart24.

### Design of the nanoengine

The two 60 nm long origami-arms are formed by 18 helix bundles (18hb) that are arranged in a honeycomb structure to form rigid and straight arms (**Fig. 1a,** for exact sequences and CadNano map see **Suppl. Dataset S1 and Suppl. Dataset S2**). They are connected by six 28 nm long dsDNA helices arranged as a 12 nm wide sheet that can be bent to serve as the compliant leaf-spring, as shown in previous work^20,25^, and by six single-stranded (ss) DNA strands that ensure the formation of the bent V-shape by being overall shorter than the double-stranded compliant area (**Fig. 1 a-e**; **Suppl. Fig. S1a**). A 154 nucleotide (nt) long transcribable dsDNA template strand (**Fig. 1 a,c**, grey; **Suppl. Fig. S1b**) spans the origami-arms and is firmly connected to each of them at 30 nm distance (**Fig. 1a**; **Fig. 1e**, red dots) from the leaf-spring ends. One of the origami-arms contains a sequence with a 5’-chloroalkyl group attached near the template dsDNA (**Fig. 1e**, yellow dot; **Suppl. Fig. S1c**), to which a HaloTag-T7RNAP fusion protein (**Fig. 1a-c**, orange-blue; **Suppl. Fig. S1b,d,e**) couples covalently at its HaloTag^26^ (HT) subunit (**Suppl. Fig. S1f**). The dsDNA template contains a T7 RNA polymerase promoter region (yellow, **Suppl. Fig. S1a,g**) and a sequence that, once transcribed, binds to a molecular beacon (green, **Suppl. Fig. S1a,g**) to monitor the amount of RNA generated during transcription. The proximity of the HT-attachment site and the dsDNA template (**Fig. 1e**, yellow and red dots) helps the T7RNAP bind to the T7 promotor sequence in the template (**Suppl. Fig. S1a,b**, yellow) and facilitates initiation of transcription in the presence of NTPs until it reaches the terminator sequence at the opposite end of the dsDNA template (**Suppl. Fig. S1a, b**, red). Moreover, the opposing origami-arm contains four biotinylated sequences that protrude the arm at the outer part of the structure to which four streptavidin proteins can be attached to unambiguously identify each arm by atomic force microscopy (AFM, **Fig. 1d,f**) or transmission electron microscopy (TEM, **Fig. 1g**). The protruding biotin residues also can potentially serve as attachment points to anchor the NE on surfaces or to other origamis. These features were designed so as to operate the NE autonomously and continuously in a rhythmic opening/closing cycle (**Fig. 1h**, **Suppl. Movie 1**) starting from the open equilibrium conformation of the structure, in which the T7-promotor is bound by the HT-T7RNAP to begin transcription (**Fig. 1h**, **1**). The pulling of the template DNA through the immobilized T7RNAP closes the origami structure while building up spring tension in the compliant segment of the structure that generates a counteracting force (**Fig. 1h**, **2**). Once the terminator sequence has been reached, the polymerase releases the template dsDNA and the structure opens to its equilibrium conformation (**Fig. 1h**, **3**) and is ready to begin a new cycle.

**Figure 1.**
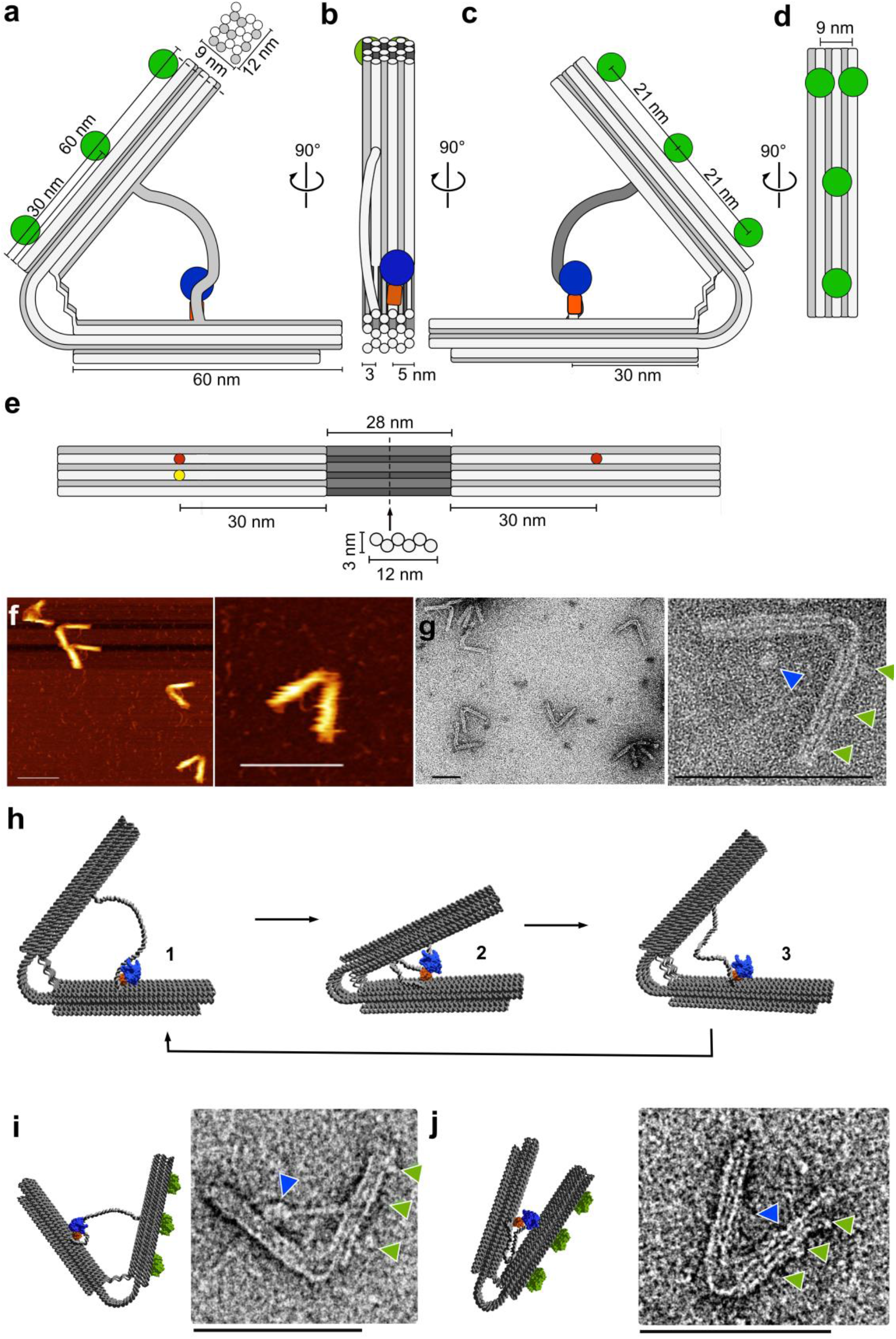
Design and dimensions of the DNA leaf-spring nanoengine. (**a**) Cross-section of the leaf-spring NE. Dimensions of the stiff origami-arms. Green circles: attachment sites for streptavidin binding; blue circle: T7RNAP-part of the HT-T7RNAP fusion protein. Top panel: arrangement and dimensions of the 18-helix bundle that forms the origami-arms, (**b**) cross section of the 90° left turn of the cross section shown in (**a**); orange: HaloTag (HT), blue: T7RNAP, (**c**) cross section and dimensions of the 90° left turn of the cross section shown in (**b**), (**d**) cross section and dimensions of the 90° left turn of the cross section shown in (**c**), (**e**) cross section and dimensions of the origami-arms that flank the 28 nm long leaf-spring helices (dark grey) that are arranged in a honeycomb structure (bottom panel), red dots: attachment sites of the dsDNA template strand, yellow dot: attachment site of the HT-T7RNAP. (**f**) Characterization by Atomic force microscopy (AFM) of the leaf-spring NE. Overview (left) and detailed image (right) of the NEs in alternating contact mode in air on a poly-L-ornithine-functionalized mica surface, (**g**) Transmission electron microscopy (TEM) of the NE in negative staining. Overview (left) and detailed image (right) of the NEs. Green arrows: streptavidin molecules bound to biotin-modified staples protruding from one of the origami-arms opposite to the location of the HT-T7RNAP fusion protein (blue arrow), (**h**) Full opening and closing cycle of the compliant mechanical structure. **1** in the open structure the dsDNA template is bound by the immobilized HT-T7RNAP fusion protein and transcription begins. **2** upon transcription, HT-T7RNAP pulls the opposing origami-arm towards itself forcing the structure to close. **3** When the terminator sequence is reached, the T7RNAP releases the dsDNA template linker, which causes the structure to snap open to its equilibrium conformation. The T7RNAP can initiate the next closing cycle. (**i**) Example of the NE engaged in transcription. Blue arrow: HT-T7RNAP, green arrows: streptavidine, (**j**) Example of the NE engaged in transcription. Blue arrow: HT-T7RNAP, green arrows: streptavidine. All scale bars: 100 nm.

The purity of the origami was analysed by gel electrophoresis (**Suppl. Fig. S2a**,**b**) and the structural integrity of NEs was confirmed by AFM (**Fig. 1f**), and TEM (**Fig. 1g**). In the AFM images of NEs that are not engaged in transcription the dsDNA template is clearly visible in between the origami-arms. The TEM images also show the HT-T7RNAP (**Fig, 1g**, right panel, blue arrow) and the three streptavidin-tags (green arrows). AFM distance measurements by height profiling confirm the designed distance of 21 nm between the streptavidin tags and the height of the origami-arms, respectively (**Suppl. Fig. S2c-e**). We also analysed NEs during transcription by TEM (**Fig. 1i,j**). The TEM image (**Fig. 1i**, right) shows that the DNA template strand exists in a strained state during the initial phase of transcription, as shown in the 3D model (left). Examples of NEs in the final phase of transcription show the origami-arms in a “closed” state (**Fig. 1j**, right), as shown in the 3D model (left).

### Functional characterization of the flat-spring nanoengine

To investigate the function of the NE we quantified the transcribed RNA by means of the MB-binding sequence encoded in the dsDNA template strand (**Fig. 2**). The fluorescence signal of the MB is expected to increase over time whenever one of the excess MB molecules hybridizes to an RNA molecule generated during transcription (**Suppl. Fig. S2f-h**). We generated a calibration curve by incubating known amounts of a complentary ODN with the MB to estimate the amount of RNA that is generated during the transcription. The importance of the HT attachment of the HT-T7RNAP fusion was measured by means of various constructs in direct relation to the transcriptional rate of the entire NE (**Fig. 2a**). In absence of the dsDNA template, no increase of fluorescence is observed, but a decrease of fluorescence due to photobleaching of the residual background fluorescence (**Fig. 2a**, column 1). The rate of transcription of the dsDNA template alone with the HT-T7RNAP shows the transcription efficiency in an intermolecular state (column 2). In a related control for intermolecular transcription, we used the NE that lacks the chloroalkane attachment site, which prevents the HT-T7RNAP from covalently attaching to the origami (column 3). The rate of transcription is comparable to that shown in column 2, indicating that there is no potential sterical hindrance by the origami and the attachment of the dsDNA template. The fully assembled NE (column 4) performs at an approximately 5 times higher transcriptional rate than the negative controls (columns 2, 3). We estimated the amount of transcript that is produces in bulk experiments and we obtained that each structure produces 2.3 ± 0.8 (n = 6, Error S.D.) transcripts each minute. These results indicate the importance of the covalent attachment of the HT-T7RNAP to the NE. The higher efficiency of transcription can be explained by proximity effects and a higher local concentration of polymerase and promotor.

**Figure 2.**
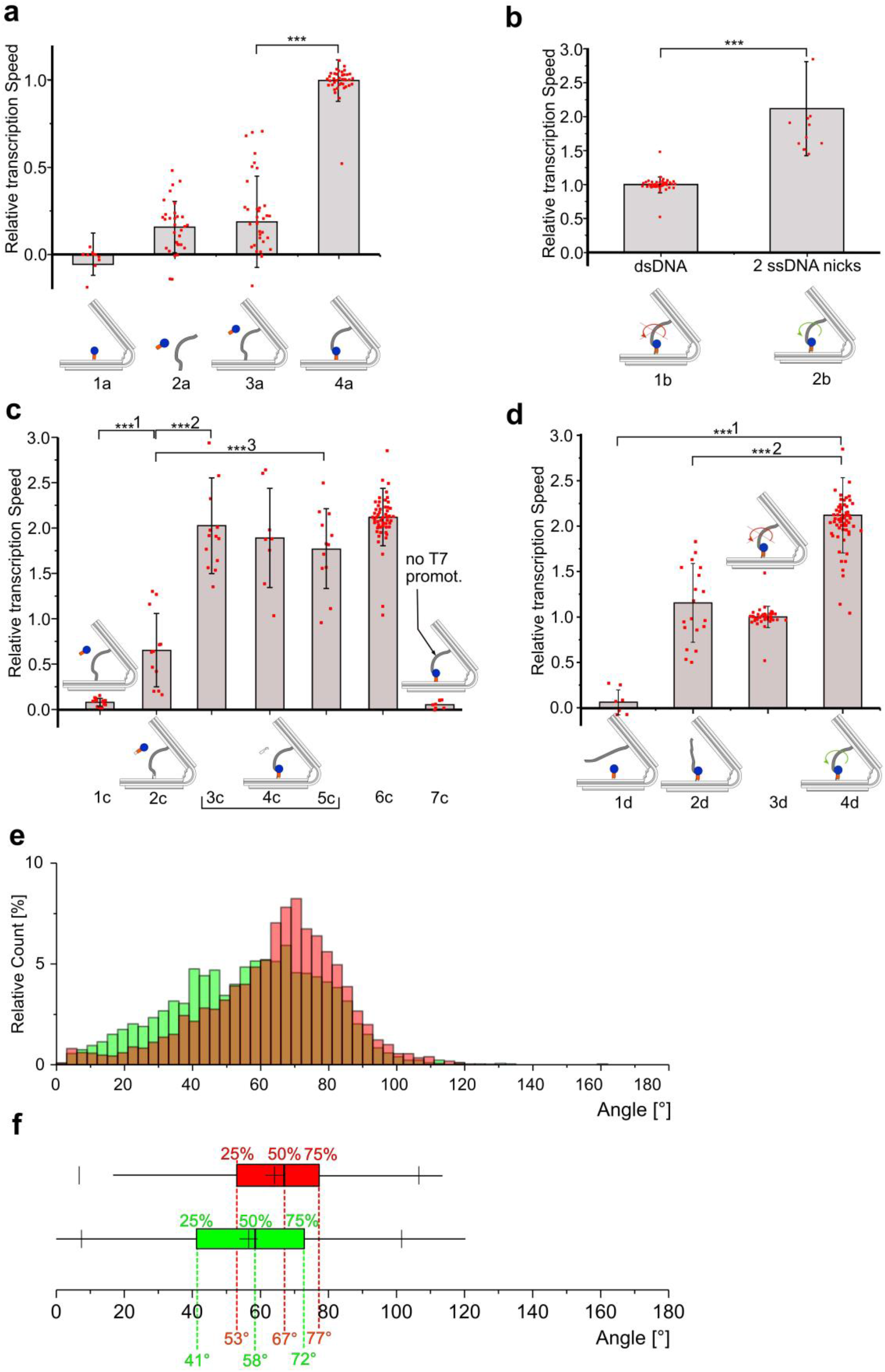
Characterization of the structures by performing bulk transcription experiments. Transcriptional rates were measured based on fluorescence increase due to MB-binding to the transcribed RNA. The transcription rates are obtained from the slope of the linear fit of the linear growth phase of the transcription curves (**Suppl. Fig. S2g-h**). All rates relate to the transcription rate of the NE, (**a**) column 1a: NE that lacks the dsDNA template, column 2a: intermolecular transcription rate from the dsDNA template alone incubated with HT-T7RNAP, column 3a: intermolecular transcription rate from the NE that lacks the chloroalkane linker, column 4a: NE. Error bars: S.D., n ≥14, ***: p = 3.0×10^-23^, (**b**) Comparison of the transcriptional rates of NE (column 1b) with nNE (column 2b). Error bars: S.D., n≥14, ***: p = 3.9×10^-5^, (**c**) column 1c: nNE that lacks the chloroalkane linker, column 2c: nNE lacking covalently bound HT-T7RNAP in presence of the HT-T7RNAP that was preincubated with 1 equivalent of the chloroalkane linker, columns 3c-5c: nNE in presence of 1 (3c), 2 (4c), and 5 (5c) equivalents of the chloroalkane linker, column 6: nNE. Error bars: S.D., n ≥8, ***1: p = 0.0005, ***2: p = 9.6×10^-8^, ***3: p = 1.6×10^-6^, column 7: nNE lacking the promotor region. Error bars: S.D., n≥7, (**d**) comparison of transcriptional rates of constructs with different attachment of the dsDNA to the origami. Column 1d: nNE in which the dsDNA template is not connected next to the HT-T7RNAP. The template has a nick at that single attachment site, column 2d: nNE in which the dsDNA template is connected next to the HT-T7RNAP. The template has a nick at that single attachment site, column 3d: NE, column 4d: nNE. Error bars: S.D., n ≥ 8, ***1: p = 1.6×10^-21^, ***2: p = 7.8×10^-10^, (**e**) Comparison of the angle distribution under the condition of transcription and no transcription. Red columns: distribution of angles in nanoengines deposited on TEM grids that did not undergo transcription (n = 5135). Green columns: distribution of angles in nanoengines that underwent transcription deposited on TEM grids (n=3266). The angles were measured using the angle measurement tool of the software imageJ (**Suppl. Dataset S3**, see also **Suppl. Fig. S3d** for a representative sample TEM image without and with indicated angles). Percentages of relative counts were calculated by dividing the counts of each bar (bin width 3°) by the total number of data in the population, (**f**) box-plot of angle distributions for the NEs not undergoing transcription (red) and the NEs that are engaged in transcription (green). Angles for the first (Q1; 25%) quartiles, the medians (Q2; 50%) and the third (Q3; 75%) quartiles are indicated by dashed lines. Whiskers: 1.5-time interquartile range. Thin vertical lines: 1 % and 99 % percentile of the distribution. Cross in box: average angle.

During transcription the T7RNAP needs to unwind the dsDNA template, which, due to its anchoring at the two origami-arms, will accumulate torsional stress as transcription proceeds. Only upon release of the polymerase at the termination signals this torsional stress can be released to build up again in the next transcription cycle, which is expected to slow the rate of transcription. The accumulation of torsional stress can be counteracted by introducing two single stranded nicks in the dsDNA strands close to the points of connection of the template to the origami (**Suppl. Fig. S2i**, red arrows). This slight structural modification permits rotation along the axle of the dsDNA template without any accumulation of torsional stress. Accordingly, we observe a two-fold increase in the rate of transcription for the NE construct containing the two nicks (nicked NE, nNE) compared to the fully double stranded sequence (**Fig. 2b, Suppl. Fig. S2iα**). To test whether changes in the compliant hinge region also influence the transcription rate, we removed two staples from the flat spring region to create two “holes” in the double stranded origami fabric, leaving only two helix strands continuously double stranded and the others partially single stranded. (**Suppl. Fig. S2j**). We hypothesized that these changes (nNEsoft) should decrease the resistance of the hinge region to the closure of the origami structure, which may increase the rate of transcription. However, we observed a comparable transcription rate of nNEsoft and nNE.

Using the nNE, we performed further control experiments for functional characterization (**Fig. 2c**). As seen before for the NE, a nNE lacking the chloroalkane linker is virtually inactive (**Fig. 2c**, column 1). Preincubation of the HT-T7RNAP with one equivalent of the chloroalkane linker to saturate the HT binding site and to prevent its covalent attachment to the attachment site in the nNE resulted in a relative transcription rate of 0.7 (column 2). However, the presence of increasing concentrations of 1, 2, and 5 equivalents of free chloroalkane linker to an assembled nNE with attached HT-T7RNAP only minimally influenced its relative transcription rate (columns 3-6), as expected. A version of nNE in which the template dsDNA lacked the promotor region (**Suppl. Fig. S2iβ**) was virtually inactive (**Fig. 2c**, column 7).

Finally, to explore the influence of the firm attachment of the dsDNA template on the relative transcription rate, we tested versions of the nNE in which the template was attached only to one of the two origami-arms (**Fig. 2d, Suppl. Fig. S2iγ,δ**). When the dsDNA template was attached only to origami-arm located opposite to the HT-T7RNAP anchored origami-arm virtually no transcription was detected (**Fig. 2d**, column 1). The most likely explanation for this observation is that the free end of the dsDNA template experiences a higher degree of freedom and electrostatic repulsion and is thus pushed away from the origami making it more difficult for the anchored polymerase to bind to the promoter region and start transcription (**Suppl. Fig. S2k**). Conversely, attaching the dsDNA template only to the site next to the HT-T7RNAP in the corresponding origami-arm resulted in a relative transcription rate of 1.4 (column 2), which is higher than that observed for the NE (column 3) but considerably lower than that of the nNE (column 4). The constructs used in columns 1 and 2 contained a single-stranded nick at the respective site of attachment. The reduced rate of construct 2d in relation to the nNE is in accordance with the notion that the higher degrees of freedom of the transcribable dsDNA and the missing “reset” function of the nNE on the dsDNA template render construct 2d less efficient than the nNE. This result suggests that the active opening of the origami in the nNE results in a straightened dsDNA template, which places the promoter region in an ideal position for the polymerase to start a new transcription cycle faster. The higher rate of construct 2d compared to the NE can be explained by the lack of torsional stress that accumulates in the transcribable strand during transcription.

Using the thousands of nNE structures displayed in the TEM images, we statistically compared the distribution of angles that the two origami-arms form in nNEs that did not undergo transcription with those that did (**SI Methods**, **Suppl. Dataset S1, Suppl. Fig. S3d**). Due to the transitional nature of the opening and closing process, in bulk experiments like these there will always be a distribution of angles over a fairly wide range, but statistically we expected to observe a broader angle distribution with a higher proportion of acute angles in the nNE samples that are engaged in transcription compared to when no transcription was occurring. The no transcription sample (**Fig. 2e**, red columns) shows a skewed distribution of angles that constantly ascends from 20° angles to reach a maximum at 70°; angles below 20° are evenly distributed at a low frequency. Above 70° the angle distribution descends rapidly towards zero indicating that a stretching of the structure above a certain angle is highly unfavoured. The transcription sample (**Fig. 2e**, green columns) exhibits a wider and flattened distribution of angle populations compared to the no-transcription curve and the median angle shifts towards more acute angles. In the angle range between 60° and 100° a considerably lower population is observed compared to the population found in the no-transcription control, whereas between 10° and 60° the population in the transcription-sample is higher. This shift towards acute angles is statistically highly significant (p=10^-57^) and demonstrates that a new population of structures is generated during the transcription in which the origami-arms form smaller angles.

Furthermore, we graphically depict the differences in the angle distributions between the two nNE samples with a boxplot showing the first (25%, Q1), second (50%, Q2, median) and third (75%, Q3) quartiles (**Fig. 2f**). The no transcription data (red box) show a narrower angle distribution than the transcription data (green box) indicating that during transcription more structures spend transitionally more time at small angles and less time in an “open” equilibrium conformation than under no-transcription conditions. The average angle for the nNE in absence of transcription is 64° ± 20° while the median resides at 67°. The Q1 is at 53°, the median at 67°, and Q3 at 77°. In nNEs that underwent transcription these values shift towards smaller angles: Q1 shifts by −12° to 41°, median shifts by −9° to 58°, Q3 shifts by −5° to 72°, and the average angle shifts by −7° to 57° ± 22°. The box for the transcribed nNEs is broader, ranging from 41° to 72° spanning 29° than the box representing the non-transcribed nNEs, which ranges from 53 to 77°, spanning 24°, which confirms a change of preferred angles towards smaller ones during transcription. The lack of a defined narrow peak in the angle distribution of nNE during transcription also indicated that the system is highly dynamic with a lack of one particular equilibrium conformation. The increase of the population of nNEs that exhibit more acute angles under transcription versus non-transcription conditions is highly significant and in accordance with the mechanism that combines the mechanical motion generated by the leaf-spring and the pulling of the immobilized HT-T7RNAP at the dsDNA template to move the two origami-arms towards each other. Taken together, the transcription experiments show that both NE and nNE function quite efficiently and behave exactly as expected.

### Molecular dynamics simulations

Coarse-grained simulations using the oxDNA model^27–30^ of different NE designs were performed to further characterize the impact of our design choices on mechanical properties (**Fig. 3a**). In addition to verifying agreement with experimental results, there were three questions that we explored using the higher degree of resolution and control afforded by molecular dynamics: (1) the relative rate of opening and closing the structure, (2) The contribution of the ds bridge to the re-opening rate, and (3) the effect of scaffold sequence across the single-stranded regions of the flexure. Notably, the simulations showed that the largest influence on mechanical properties came from transient, sequence-dependent secondary structures formed by the single-stranded domains in the flexure. Disallowing formation of these secondary structures (“no structure”: all structures labelled as NS in **Fig. 3b**) increased both the equilibrium angle and the stiffness of the (n)NE (**Fig. 3b**). Conversely, allowing the formation of secondary structures reduces the equilibrium angle to roughly the same value, irrespective of variations in the transcribable sequence (NE_NB, NE, and nNE in **Fig 3b**). This effect is particularly pronounced in the NE-structure lacking the transcribable DNA sequence (“no-bridge”: labelled as NB in **Fig. 3b**), where the difference in equilibrium angle between NE_NB and NE_NS_NB is 25°. This result suggests that the secondary structures in the hinge region are the largest determinant of equilibrium angle and stiffness (**Suppl. Fig. S4a**); however, the presence of the transcribable sequence imposes a hard upper limit on the opening angle of the structure.

**Figure 3.**
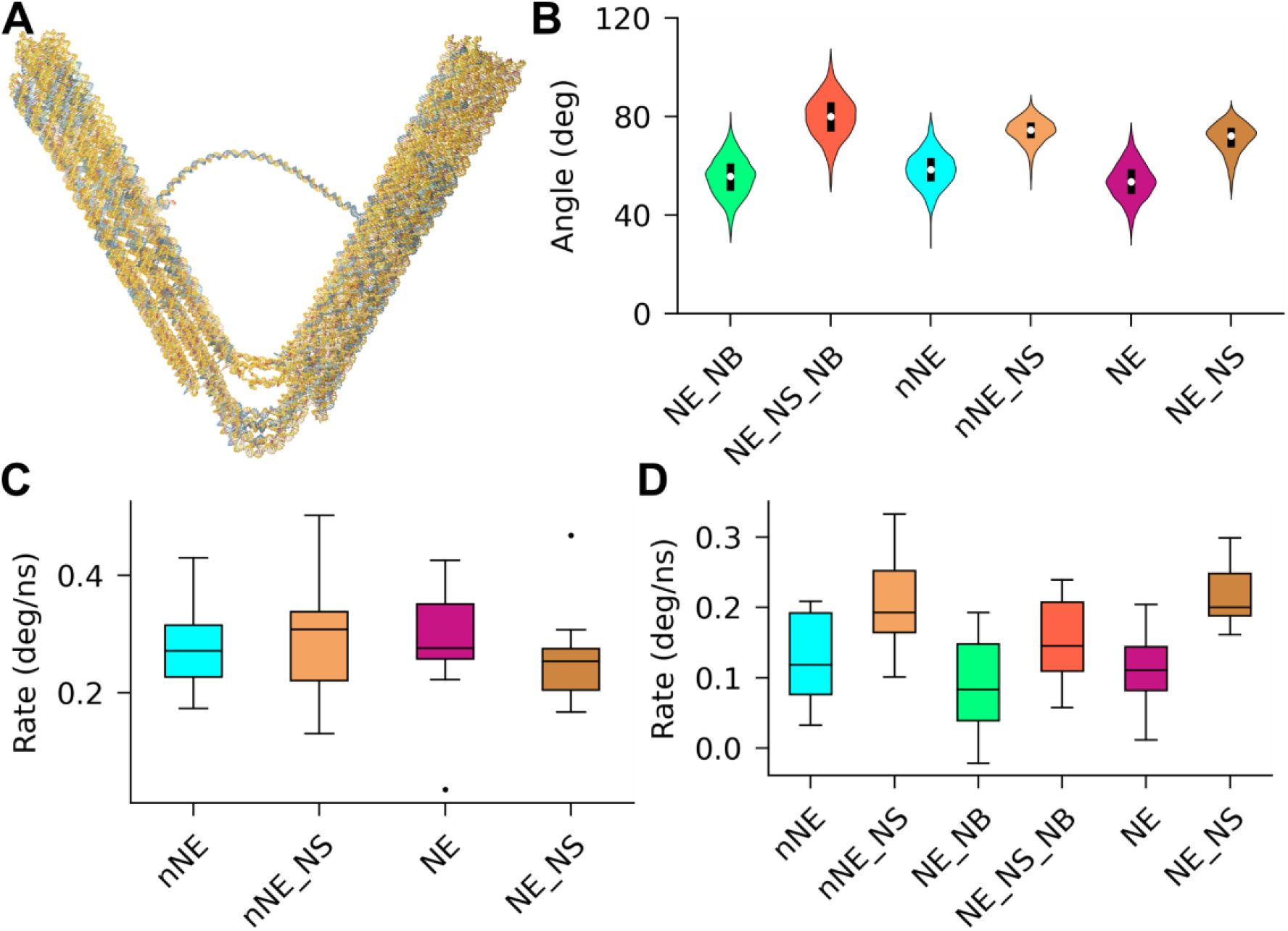
Coarse-grained simulation of NE designs. oxDNA simulations were performed to determine dynamic structural properties of the NE. (**a**) Mean structure of an equilibrium sampling simulation of nNE represented in oxDNA, (**b**) Equilibrium angle distribution of six designs during oxDNA simulation. NE - nano engine; nNE - nicked nano engine; NS - no structure, where base pairing was turned off for single stranded regions of the flexure to isolate the effect of secondary structures forming in the flexure; NB - no transcribable dsDNA bridge; here the transcribable dsDNA bridge was deleted using oxView’s editing tools, to determine its effect on equilibrium and opening simulations, (**c**) Simulation-determined closing rates from pulling simulations, (**d**) simulation-determined opening rates from relaxation simulations where the forces from the pulling simulations are released and the structure is allowed to open again.

In the absence of the transcribable dsDNA and secondary structures in the hinge region, the DNA leaf spring can stretch out and reach an equilibrium angle of 80° with a standard deviation of 9°. When the transcribable sequence is introduced, NE_NS and nNE_NS have comparable equilibrium angles of 71 ±6° and 74 ±5°, respectively, both with a negatively skewed distribution caused by the upper limit imposed by the transcribable sequence. When secondary structures are permitted, the scenario changes drastically: now NE_NB, nNE, and NE have highly comparable equilibrium angles of 55° ±8°, 58° ±7°, and 54° ±7° respectively (**Fig. 3b**), implying that the effect of the transcribable dsDNA on the equilibrium angle is negligible. These results suggest that the hairpins formed by the single stranded sequences decrease the length and flexibility of the hinge region, influencing both the overall shape and rigidity of the origami. The structures in the hinge region are only slightly affected by temperature changes. Increasing the simulation temperature from 23 °C to 37 °C only marginally increases the angles observed in the simulated structures. (**Suppl. Fig. S4b**).

To test whether the predicted effects of the simulations can be confirmed by experimental results, we analysed the angle distribution using TEM images of a construct lacking the transcribable dsDNA (NE-NB) and compared it with that of NE (**Suppl. Fig. S4c**, **d**). We observed that the two populations have the exact same distribution (p = 0.6), suggesting that the secondary structures predicted by the simulations are indeed present in experimental conditions and have an impact on the conformational freedom of the structurers. Notably, the angle was systematically lower in the simulation compared to the experimental measurements. A possible reason for the difference is that the experimental angles were measured from TEM images of surface deposited structures, which could have caused slight deformation of the structure due to the adherence to the TEM grid. In the 3D simulations we observe out-of-plane twisting in the structures (**Suppl. Fig. S4e**). It is likely that deposition of structures on the surface compensates some of this out-of-plane twist by opening the structure slightly wider.

To sample the behaviour of the NE under tension, mimicking the action of T7RNAP, pulling and release simulations were performed by applying a constant force comparable to that exerted by the T7RNAP (16 pN) between the nucleotide covalently linked to the T7RNAP in experiments and the first nucleotide of the terminator sequence. In general, closing was faster than opening with rates of 0.28 ±0.07 °/ns vs. 0.13 ±0.06 °/ns for nNE and 0.28 ±0.10 °/ns vs. 0.11 ±0.05 °/ns for NE. These results are in accordance with what one would expect if we take into account that closing is a driven process while opening relies on releasing strain built up during the closing process and a component of Brownian motion. It also explains our experimental observation that nNEsoft showed a similar transcription rate as nNE (**Suppl. Fig. S2j**). Interestingly, when secondary structures are prohibited the closing rate for nNE-NS was 0.29 ±0.10 °/ns and the opening rate was 0.21 ±0.07 °/ns. For NE-NS, the closing rate was 0.26 ±0.08 °/ns and the opening rate was 0.22 ±0.05 °/ns (**Fig. 3d**). The similarity of the closing rates while doubling the opening rates when secondary structures are disallowed suggests that the secondary structures have little effect on the driven closing process, however they stabilize the closed state and impede the stochastic opening process. The transcribable sequence has a smaller impact on opening rates. In simulations where that region was deleted from the closed state and allowed to re-open, the rate was slightly slower than the corresponding complete structure (0.08 ±0.07 °/ns for NE_NB and 0.15 ±0.06 °/ns for NE_NS_NB). No significant difference was observed between the rates of pulling and opening when NE (with the entirely transcribable dsDNA) and nNE (where the transcribable dsDNA contained two nicks) simulations initiated from the same starting configuration (**Fig. 3c, d**). The simulation results hence suggest that the experimentally observed difference in closing and opening rates between NE and nNE can be attributed to supercoiling caused by the activity of the polymerase rather than to the mechanical properties conferred by the transcribable dsDNA to the NE itself.

The effect of temperature on the dynamic simulations was similar to the equilibrium simulations. The rates of opening and closing were slightly increased and the variance of replicates of closing and opening simulations increased at 37 °C, as would be expected due to the increase in the accessible phase space (**Suppl. Fig. S4f, g**).

To get a better insight into the observed phenomenon of a reduced transcription rate when the transcribable dsDNA sequence is anchored only next to the polymerase (**Fig. 2d**) we extracted the amount of time the promoter region spends next to the polymerase in the case when the dsDNA strand is fully anchored at both ends to the origami and when it is only attached next to the polymerase. As the T7RNAP is not explicitly represented in the simulations, an approximation of accessibility was made based on distance. The overall geometry of T7RNAP is roughly that of a sphere 10.35 nm in diameter31. Thus, to a first order approximation, transcription can only be initiated when the distance between the polymerase attachment site and promoter sequence are within 10.35 nm proximity (this approximation ignores the specific orientation required to initiate transcription). The simulations showed that in the nNE the promoter region spends 54.9 ±0.01% of the time in the 10.35 nm radius but this time is reduced to 45.6 ±0.01% when the dsDNA strand is only anchored next to the polymerase indicating how the higher number of degrees of freedom in the latter case reduces the chances of the promoter region to be in a favourable position to obtain efficient transcription. (**Suppl. Fig. S4h**).

### Driver-follower experiments

An important task of any engine that actively performs work is the ability to be coupled to passive moving parts for transmission of force or motion. Nature has found vast solutions for the transmission of force, primarily by myocytes or adherent cells, but examples demonstrating nanomechanical force and motion transmission by synthetic machines are scarce and the motion occurs mostly stochastic^32–35^. To demonstrate that nNE can act as non-stochastic, autonomous mechanical ‘driver’ (D, **Fig. 4a**) and actively transmit its force to a passive part that follows its motion, we coupled nNE to a passive ‘follower’ (F, **Fig. 4b**). We added five unique ssDNA overhangs protruding axially from the extremities of each origami arm of D and F. Each of the ten sequences in D allow for the hybridization of only the complementary 10 sequences on the F unit to form the rhomboid-shaped hetero dimer D-F (**Fig. 4c-e**); formation of homodimers D-D or F-F is not possible due to the non complementarity of the sequences (**Suppl. Fig. S5a, b**). To strengthen the connection between D and F, three LNA modifications^36^ were included into two of the overhanging sequences on each D-arm that connect to F (**Suppl. Fig. S5b**, red).

**Figure 4.**
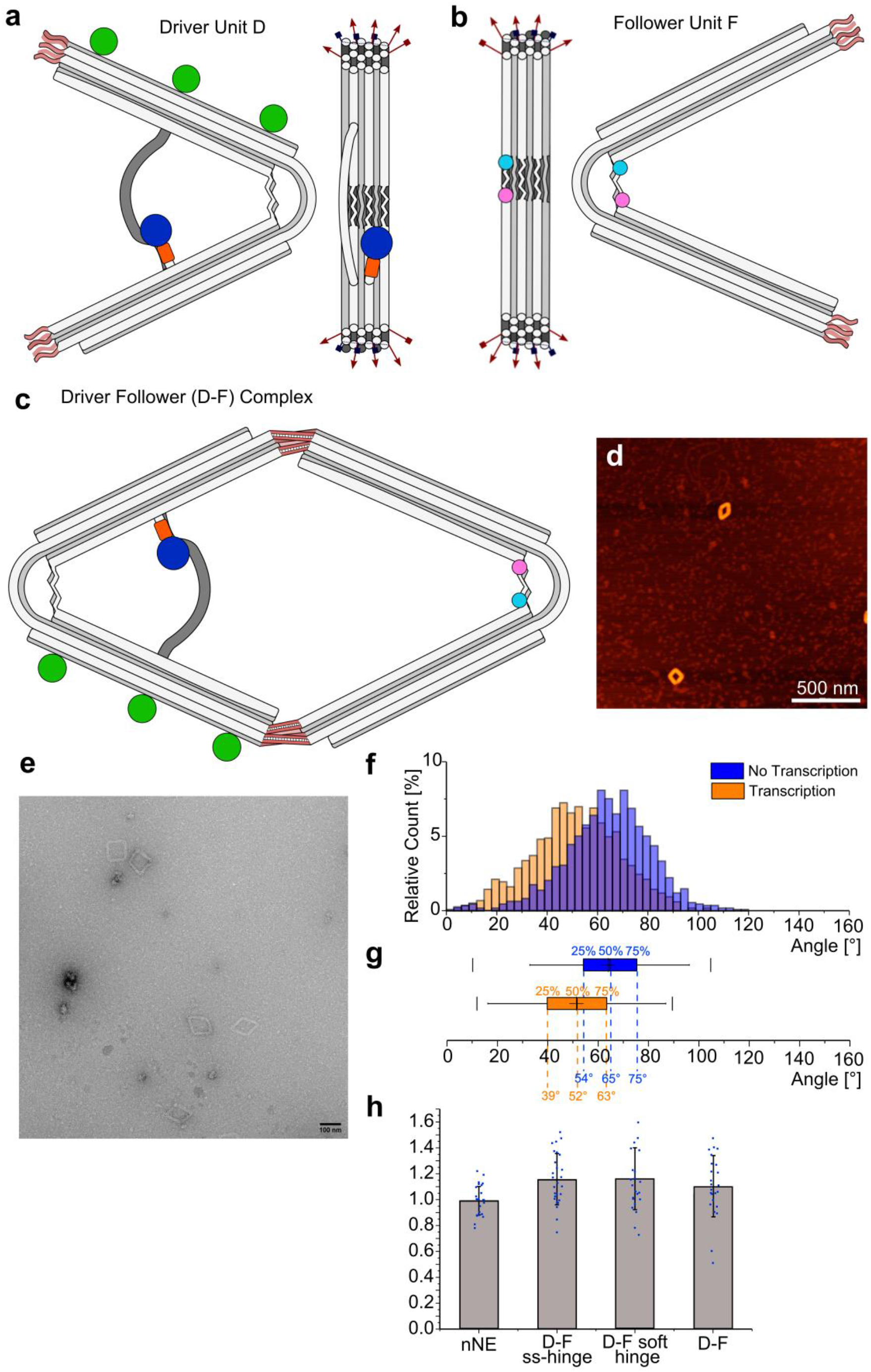
nNE drives a passive follower unit. (**a**) Schematic representation of the nNE driver D and the passive follower unit F in (**b**). The structures are modified to have ssDNA overhangs protruding from the stiff origami arms. Five ssDNA overhangs (shown as red arrows) were introduced on each arm for a total of 10 unique overhangs on each origami. The sequences are designed to allow the connection of only the active structure to the passive structure. Biotin modifications are introduced only on the driver unit and are shown as green dots. A Cy3 and Cy5 FRET pair in F are shown with a pink and cyan dot, (**c**) the assembly of the D-F heterodimer is achieved by equimolarly combining the two parts of the system and allowing a thermal annealing overnight. After 4.5 h of transcription experiments at 37 °C the integrity of the system is confimed by AFM (**d**) size bar: 500 nm, and TEM (**e**) size bar: 100 nm, (**f**) Analysis of the angle distribution from TEM images of the D-F dimer complex for samples that did not (n=1074, blue) and did undergo transcription (n=1190, orange). The distribution was plotted as bar graphs with a bin with of 3°, the count of each bar was divided by the total number of counts, and displayed as the relative count percentages. The distribution significantly shifts towards more acute angles under transcription (p = 8 x 10^-64^), indicating the formation of structures with smaller angles during transcription; (**g**) Boxplot representation of distributions. Blue: boxplot of D-F without transcription, orange: box plot of D-F with transcription. The boxes represent the distribution at first quartile (Q1; 25%), second quartile or median (Q2; 50%) and the third quartile (Q3; 75%) with dashed lines indicating the angular value in the corresponding color. The whiskers represent 1.5-times the box size while the thin vertical lines represent the 1% and the 99% percentile of the distribution. The thin cross indicates the average angle of the distribution. Average angles of distribution: D-F in absence of transcription 64°±17° (S.D. error), D-F with transcription 51° ±17°, (**h**) The transcription speed of different D-F complexes was measured and compared to the transcription speed of D (nNE) alone (n = 27). Transcription speeds: D-F-ss-hinge 1.2 ±0.2 (n = 27, p = 0.001), D-F-soft-hinge 1.2 ±0.2 (n = 18, p = 0.01), D-F complex 1.1 ±0.2 (n = 24; p = 0.06). Bargraph: average value with error S.D.

As before for nNE, a large set of TEM images of only the rhomboidal D-F dimers (**Suppl. Figure S5c**) with and without transcription was analysed to obtain angle distributions under both conditions (**Fig. 4f**). Box plots of the distribution (**Fig. 4g**) show that the median (Q2) shifts from 65° in absence of transcription (blue) to 52° in presence (orange). The quartile distribution Q1 shifts from 54° (no transcription, blue) to 39° (transcription, orange), and Q3 shifts from 75° (no transcription, blue) to 63° (transcription, orange). Moreover, the formation of the D-F complex apparently leads to slightly narrower, less skewed, and more symmetric angle distributions under both conditions (**Suppl. Fig. S5d**, blue, orange) as compared to NE (red, green) suggesting that the connection of D and F has a stabilizing effect on the angle. These data demonstrate that transcribing D-F structures exhibit more acute angles on average than non-transcribing ones and indicate that the D-bound F unit follows the motion imposed by D.

This trend becomes even more apparent in case of a D-F complex in which the F unit has a completely single stranded flexure region (F-ss-hinge, **Suppl. Fig. S5f, g**). In the ‘ss-hinge’-design the interaction of dsDNA flat spring and ssDNA tension sequences is absent. Consequently, F does not assume a defined angle, which is reflected by a broad angle distribution median of 113° (**Suppl. Fig. S5h, i**, olive, F-ss-hinge). However, when bound to D, the median shifts towards a more acute median angle of 71° (**Suppl. Fig. S5h, i**, wine, D-F-ss-hinge no transcription) under no transcription, and a median of 56° under transcription conditions (**Suppl. Fig. S5h, i**, green, D-F-ss-hinge transcription), comparable with the medians obtained for NE. The boxplots also show that the angle distributions become considerably narrower in the D-F-ss-hinge complex compared to F-ss-hinge: under no transcription the difference beween Q3 and Q1 in D-F-ss-hinge measures only 23° while in the F-ss-hinge it measures 54° (**Suppl. Fig. S5i**).

When the D unit was coupled to an F-derivative with a soft hinge, (F-soft-hinge; **Suppl. Fig. S5j**) in which two staples are removed from the dsDNA flexure region, the transcription shows a behaviour that is comparable with the D-F sample. The median angle of the distribution shifts from 62° in absence of transcription (yellow, **Suppl. Fig. S5k, l**) to 50° under transcription conditions (purple).

We next measured the transcription speed of different driver follower constructs relative to the single D (or nNE) unit. D was combined with the F-ss-hinge, F-soft-hinge, or the F unit (**Fig. 4h**). Although the differences are small, we observed a significant increase of transcription rate for the D-F-ss-hinge and the D-F-soft-hinge, respectively, of 1.2 ±0.2 (p = 0.001) and 1.2 ±0.2 (p value 0.01), respectively, compared to nNE. The combination of D with F (D-F) showed no significant increase in transcription speed compared to nNE (**Fig. 4h**, nNE vs. D-F). These results indicate that the combination of the active driver with a passive follower unit influences the closing and opening speed of the dimeric system. In case of the D-F we hypothesize that the addition of a complete hinge to the dimeric complex adds further resistance to the closing and this effect is compensated just enough by the reopening spring effect to result in no significant effect on the overall transcription speed. In the case of the F-ss-hinge and the F-soft-hinge, however, the closing speed is not significantly affected due to less resistant springs but the reopening is enhanced by the higher entropic degree of spring areas resulting in a slightly increased transcription rate.

### RNA transcript analysis

To further corroborate the expected behavior of the nanoengines and the D-F-pair, we directly compared the amounts and lengths of transcribed RNA molecules produced by the various nanoengines and controls used in this study (**Fig. 5**) by polyacrylamide gel electrophoresis (PAGE). For NE, nNE, and Driver D exactly the same major transcription products are obtained that correspond to RNAs that terminated either at the first or at the second terminator sequence (**Fig. 5a**, lanes 1-3). For the nNE control in which the transcribable DNA is anchored next to the polymerase only (**Fig. 5a**, lane 4, for the construct see **Fig. S2i∂**) the major RNA product corresponds to the one that ends after the first terminator, since the DNA template sequence ends before the second terminator sequence. In the nNE control that lacks the dsDNA template no band is visible (**Fig. 5a**, lane 6). The nNE control without the chloroalkane linker that shows drastically reduced transcription rates since HT-T7RNAP is not anchored next to the transcribable sequence (**Fig. 5a**, lane 5) shows no transcribed RNA either, and the same is true for just the HaloTag-T7RNAP incubated with the free transcribable DNA template both for nNE and NE (**Fig. 5a**, lanes 7, 8). The same set of experiments was analyzed by PAGE under denaturing conditions (**Fig. 5b**). In this case the transcripts of NE, nNE, Driver, and the construct shown in **Fig. S2i∂** run at the same levels. This migration behaviour indicates that the transcripts in all four versions are of comparable length and that the differences observed under non denaturing conditions can be attributed to secondary structures that form in the produced RNA. Furtherore, the similar transcription product lengths obtained for NE, nNE, D, and for the construct shown in **Fig. S2i∂** excludes the unlikely possibility that HT-T7RNAP reads through the terminator sequences in NE, nNE, or D, and continues transcribing the staple-section that anchores the DNA-template within the origami, which would compromise the pulling force. Finally, we also analyzed two independent batches of the genuine D-F pair in the same way as D only, and found a similar RNA pattern as observed for D only (**Suppl. Fig. S6**). Taken together, these data demonstrate that NE, nNE, D, and D-F behave as designed and expected.

**Figure 5.**
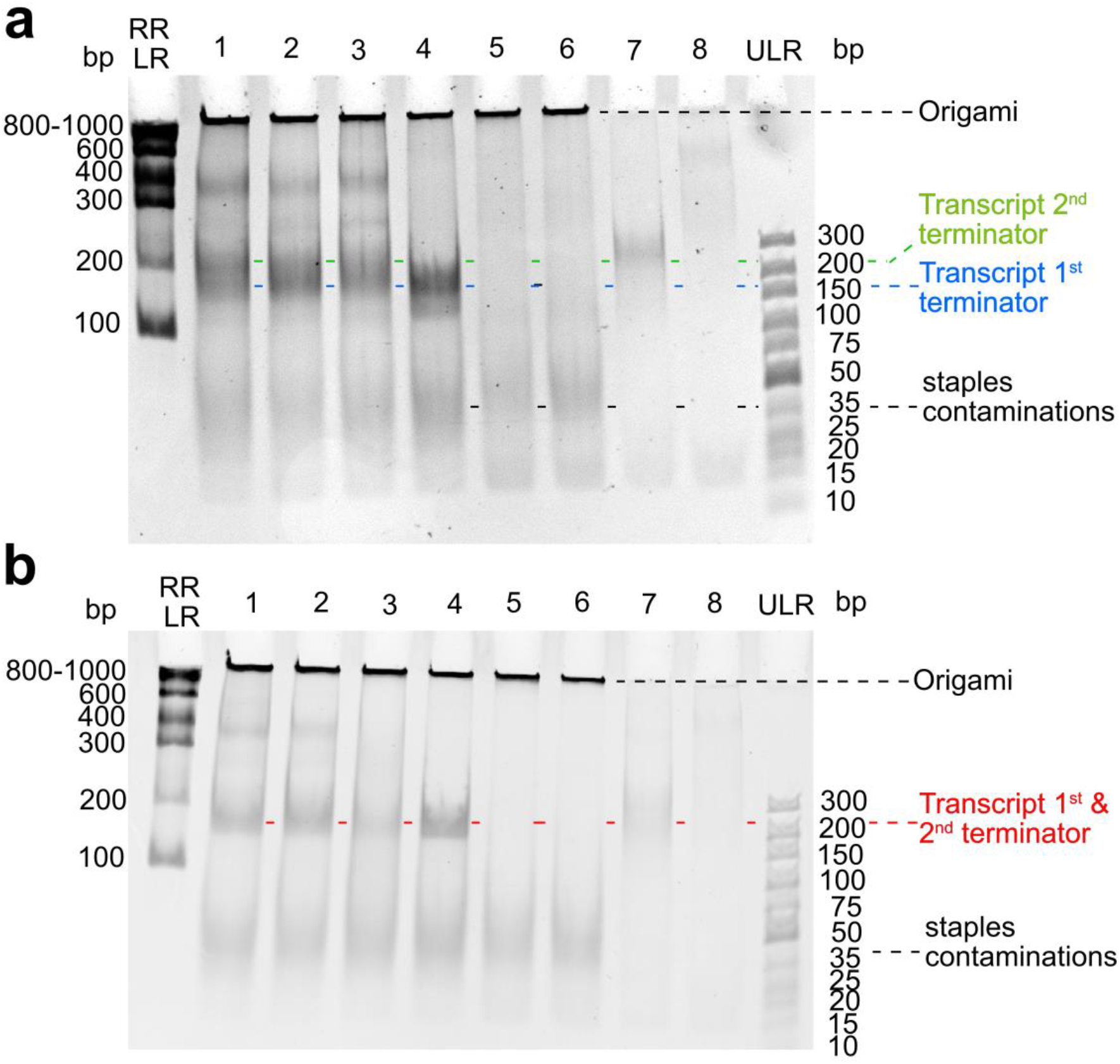
Amounts of RNA observed by polyacrylamide gel electrophoresis (PAGE) for nanoengines NE, nNE, D, and controls. (**a**) Transcription products after 4 h at 37° C (6% PAGE) for the constructs NE (lane 1); nNE (lane 2); driver D (lane 3); nNE with DNA template attached only next to T7RNAP (lane 4; see Fig. S2i*∂*); nNE without the Chloroalkane modified staple (lane 5, construct 3a in Fig. 2a); nNE without the transcribable DNA sequence (lane 6, construct 1a in Fig. 2a); transcribable DNA sequence for nNE only (lane 7); transcribable DNA sequence for the NE only (lane 8, construct 2a in Fig. 2a); RR-LR: RiboRuler Low Range ss RNA ladder; ULR: Ultra low range dsDNA ladder, (**b**) the same set of experiments run under denaturing 6% PAGE conditions. Samples were incubated in 8.4 M urea at 35 °C for 10 min before loading on the gel.

## Conclusion

We describe the bottom-up construction of a biohybrid DNA-origami-based nanomachine that performs tasks that are fundamental for every contrivance requiring automated motion: An autonomous, fuel-driven, rhythmically pulsing DNA-nanoengine that can be easily latched to any type of passive DNA-origami-based ‘follower’ entities to which it then transmits its motion and force, thus constituting a genuine driver-follower pair.

Achieving autonomous active motion in a DNA-nanostructure constitutes a success in itself, but the demonstration that nNE can act as a driver of other devices adds considerable significance. Since DNA origami technology permits the bottom-up construction of robust structures with different mechanical properties that can span from mechanically rigid to bendable and compliant structures^11, 19–20, 25^, all bearing the option of being combined into a single architecture, the versatility of mechanical power transmission by nNE is vast. Although the different passive follower structures used here are fairly simple, the prototypical design of D-F suggests that nNE should be applicable in other DNA-nanostructures to achieve considerably larger structural rearrangements as exemplified before in non-autonomous systems^37^. nNE provides an actively moving nano engine that operates fully autonomously; once the NTP-fuel is added to the system, the structure sets itself in motion, and continues pulsation over several hours without further input. It can even be envisioned introducing a switchable clutch mechanism that straightforwardly allows detaching the driver from a coupled follower and attaching it to a different one while the engine is still running. For example by controllong the hybridisation of the D-F connecting ODNs with light-switchable isomers, as has been shown for several DNA nanomachines^10, 38–42^.

By expanding the complexity of the structural features that can be easily attached to the origami by single stranded overhangs or other chemical or biochemical modification, there is potentially no limit to the range of motion or the type of motion that can be linear, rotative, pulsatile or contractile, that could be achieved by introducing the nano engine as an active core component at the heart of bigger passive nano machines, the function of which depends on downstream sequences of motion.

## Acknowledgements

We thank V. Fieberg, and D. Keppner for technical assistance, and Werner Kühlbrandt, MPI of Biophysics, Frankfurt, for providing access to electron microscopy facilities. This work was supported by the Alexander von Humboldt Foundation, the Max-Planck Society and the University of Bonn. P. Š. and E. P. acknowledge the use of the Extreme Science and Engineering Discovery Environment (XSEDE), which is supported by National Science Foundation grant number TG-BIO210009.

## Authors Contributions

M. F., Mat. C., and J. V. developed the here described concept of the T7RNAP-driven nanoengine. Mat.C. performed and designed, with M. F. and initially also with J. V., most of the included studies. M. F. supervised the research project, assisted by J. V. Mar. C. performed all TEM studies. E. P. and P. Š. carried out all molecular modelling. All authors discussed the experimental results and contributed to writing the manuscript, with Mat. C. and M. F. doing the bulk of the writing.

## Additional information

The authors declare no competing financial interests. The input files and analysis scripts for the simulation portions of this work are available at https://github.com/sulcgroup/hinges, while the full oxDNA trajectories can be found at https://drive.google.com/drive/folders/1KoocIZRPcRJ7us0q695_ya3cXNxYUn2t. Supplementary Datasets S1-S3 and Supplementary Movie S1 can be found at https://drive.google.com/drive/folders/1gp6lGHHh55W4BCZR7tnTNTAuUsph-GQQ?usp=sharing. Reprints and permission information is available online at http://…. Correspondence and requests for materials should be addressed to.

## Notes

https://github.com/sulcgroup/hinges

https://drive.google.com/drive/folders/1KoocIZRPcRJ7us0q695_ya3cXNxYUn2t

https://drive.google.com/drive/folders/1gp6lGHHh55W4BCZR7tnTNTAuUsph-GQQ?usp=sharing

### Competing Interest Statement

The authors have declared no competing interest.

## References

1. Kammerer, C.; Erbland, G.; Gisbert, Y.; Nishino, T.; Yasuhara, K.; Rapenne, G., Biomimetic and Technomimetic Single Molecular Machines. Chem Lett 2019, 48 (4), 299–308.

2. Feringa, B. L., The art of building small: from molecular switches to molecular motors. J. Org. Chem. 2007, 72 (18), 6635–52.

3. Stoddart, J. F., Mechanically Interlocked Molecules (MIMs)-Molecular Shuttles, Switches, and Machines (Nobel Lecture). Angew Chem Int Ed Engl 2017, 56 (37), 11094–11125.

4. Sauvage, J. P., From Chemical Topology to Molecular Machines (Nobel Lecture). Angew Chem Int Ed Engl 2017, 56 (37), 11080–11093.

5. Bath, J.; Turberfield, A. J., DNA nanomachines. Nat. Nanotechnol. 2007, 2 (5), 275–84.

6. Erbas-Cakmak, S.; Leigh, D. A.; McTernan, C. T.; Nussbaumer, A. L., Artificial Molecular Machines. Chem Rev 2015, 115 (18), 10081–206.

7. Feng, Y.; Ovalle, M.; Seale, J. S. W.; Lee, C. K.; Kim, D. J.; Astumian, R. D.; Stoddart, J. F., Molecular Pumps and Motors. J Am Chem Soc 2021.

8. von Delius, M.; Leigh, D. A., Walking molecules. Chem. Soc. Rev. 2011, 40 (7), 3656–76.

9. Chakraborty, K.; Veetil, A. T.; Jaffrey, S. R.; Krishnan, Y., Nucleic Acid-Based Nanodevices in Biological Imaging. Annu Rev Biochem 2016, 85, 349–73.

10. Kamiya, Y.; Asanuma, H., Light-driven DNA nanomachine with a photoresponsive molecular engine. Acc Chem Res 2014, 47 (6), 1663–72.

11. Marras, A. E.; Zhou, L.; Su, H. J.; Castro, C. E., Programmable motion of DNA origami mechanisms. Proc Natl Acad Sci U S A 2015, 112 (3), 713–8.

12. Amano, S.; Fielden, S. D. P.; Leigh, D. A., A catalysis-driven artificial molecular pump. Nature 2021, 594 (7864), 529–534.

13. Feng, L.; Qiu, Y.; Guo, Q. H.; Chen, Z.; Seale, J. S. W.; He, K.; Wu, H.; Feng, Y.; Farha, O. K.; Astumian, R. D.; Stoddart, J. F., Active mechanisorption driven by pumping cassettes. Science 2021, 374 (6572), 1215–1221.

14. Erbas-Cakmak, S.; Fielden, S. D. P.; Karaca, U.; Leigh, D. A.; McTernan, C. T.; Tetlow, D. J.; Wilson, M. R., Rotary and linear molecular motors driven by pulses of a chemical fuel. Science 2017, 358 (6361), 340–343.

15. Ragazzon, G.; Baroncini, M.; Silvi, S.; Venturi, M.; Credi, A., Light-powered autonomous and directional molecular motion of a dissipative self-assembling system. Nat Nanotechnol 2015, 10 (1), 70–5.

16. Kudernac, T.; Ruangsupapichat, N.; Parschau, M.; Macia, B.; Katsonis, N.; Harutyunyan, S. R.; Ernst, K. H.; Feringa, B. L., Electrically driven directional motion of a four-wheeled molecule on a metal surface. Nature 2011, 479 (7372), 208–11.

17. Baroncini, M.; Casimiro, L.; de Vet, C.; Groppi, J.; Silvi, S.; Credi, A., Making and Operating Molecular Machines: A Multidisciplinary Challenge. ChemistryOpen 2018, 7 (2), 169–179.

18. Wilson, M. R.; Sola, J.; Carlone, A.; Goldup, S. M.; Lebrasseur, N.; Leigh, D. A., An autonomous chemically fuelled small-molecule motor. Nature 2016, 534 (7606), 235–40.

19. DeLuca, M.; Shi, Z.; Castro, C. E.; Arya, G., Dynamic DNA nanotechnology: toward functional nanoscale devices. Nanoscale Horiz 2020, 5 (2), 182–201.

20. Zhou, L.; Marras, A. E.; Su, H. J.; Castro, C. E., DNA origami compliant nanostructures with tunable mechanical properties. ACS Nano 2014, 8 (1), 27–34.

21. Valero, J.; Famulok, M., Regeneration of Burnt Bridges on a DNA Catenane Walker. Angew Chem Int Ed Engl 2020, 59 (38), 16366–16370.

22. Valero, J.; Pal, N.; Dhakal, S.; Walter, N. G.; Famulok, M., A bio-hybrid DNA rotor-stator nanoengine that moves along predefined tracks. Nat Nanotechnol 2018, 13 (6), 496–503.

23. Yu, Z.; Centola, M.; Valero, J.; Matthies, M.; Sulc, P.; Famulok, M., A Self-Regulating DNA Rotaxane Linear Actuator Driven by Chemical Energy. J. Am. Chem. Soc. 2021, 143 (33), 13292–13298.

24. Wikipedia Leonardo’s self-propelled cart. https://en.wikipedia.org/wiki/Leonardo%27s_self-propelled_cart.

25. Shi, Z.; Castro, C. E.; Arya, G., Conformational Dynamics of Mechanically Compliant DNA Nanostructures from Coarse-Grained Molecular Dynamics Simulations. ACS Nano 2017, 11 (5), 4617–4630.

26. Los, G. V.; Encell, L. P.; McDougall, M. G.; Hartzell, D. D.; Karassina, N.; Zimprich, C.; Wood, M. G.; Learish, R.; Ohana, R. F.; Urh, M.; Simpson, D.; Mendez, J.; Zimmerman, K.; Otto, P.; Vidugiris, G.; Zhu, J.; Darzins, A.; Klaubert, D. H.; Bulleit, R. F.; Wood, K. V., HaloTag: a novel protein labeling technology for cell imaging and protein analysis. ACS Chem Biol 2008, 3 (6), 373–82.

27. Ouldridge, T. E.; Louis, A. A.; Doye, J. P. K., Structural, mechanical, and thermodynamic properties of a coarse-grained DNA model. J Chem Phys 2011, 134 (8).

28. Rovigatti, L.; Sulc, P.; Reguly, I. Z.; Romano, F., A comparison between parallelization approaches in molecular dynamics simulations on GPUs. J Comput Chem 2015, 36 (1), 1–8.

29. Snodin, B. E.; Randisi, F.; Mosayebi, M.; Sulc, P.; Schreck, J. S.; Romano, F.; Ouldridge, T. E.; Tsukanov, R.; Nir, E.; Louis, A. A.; Doye, J. P., Introducing improved structural properties and salt dependence into a coarse-grained model of DNA. J Chem Phys 2015, 142 (23), 234901.

30. Sulc, P.; Romano, F.; Ouldridge, T. E.; Rovigatti, L.; Doye, J. P. K.; Louis, A. A., Sequence-dependent thermodynamics of a coarse-grained DNA model. J Chem Phys 2012, 137 (13).

31. Durniak, K. J.; Bailey, S.; Steitz, T. A., The structure of a transcribing T7 RNA polymerase in transition from initiation to elongation. Science 2008, 322 (5901), 553.

32. Ramezani, H.; Dietz, H., Building machines with DNA molecules. Nat Rev Genet 2020, 21 (1), 5–26.

33. Yoon, J.; Eyster, T. W.; Misra, A. C.; Lahann, J., Cardiomyocyte-Driven Actuation in Biohybrid Microcylinders. Adv Mater 2015, 27 (30), 4509–4515.

34. Sagara, Y.; Karman, M.; Verde-Sesto, E.; Matsuo, K.; Kim, Y.; Tamaoki, N.; Weder, C., Rotaxanes as Mechanochromic Fluorescent Force Transducers in Polymers. J. Am. Chem.Soc. 2018, 140 (5), 1584–1587.

35. Chen, S.; Wang, Y.; Nie, T.; Bao, C.; Wang, C.; Xu, T.; Lin, Q.; Qu, D. H.; Gong, X.; Yang, Y.; Zhu, L.; Tian, H., An Artificial Molecular Shuttle Operates in Lipid Bilayers for Ion Transport. J Am Chem Soc 2018, 140 (51), 17992–17998.

36. Vester, B.; Wengel, J., LNA (locked nucleic acid): high-affinity targeting of complementary RNA and DNA. Biochemistry 2004, 43 (42), 13233–41.

37. Gerling, T.; Wagenbauer, K. F.; Neuner, A. M.; Dietz, H., Dynamic DNA devices and assemblies formed by shape-complementary, non-base pairing 3D components. Science 2015, 347 (6229), 1446–52.

38. Skugor, M.; Valero, J.; Murayama, K.; Centola, M.; Asanuma, H.; Famulok, M., Orthogonally Photocontrolled Non-Autonomous DNA Walker. Angew Chem Int Ed Engl 2019, 58 (21), 6948–6951.

39. Wang, S.; Yue, L.; Li, Z. Y.; Zhang, J.; Tian, H.; Willner, I., Light-Induced Reversible Reconfiguration of DNA-Based Constitutional Dynamic Networks: Application to Switchable Catalysis. Angew Chem Int Ed Engl 2018, 57 (27), 8105–8109.

40. Asanuma, H.; Ito, T.; Yoshida, T.; Liang, X.; Komiyama, M., Photoregulation of the Formation and Dissociation of a DNA Duplex by Using the cis-trans Isomerization of Azobenzene. Angew Chem Int Ed Engl 1999, 38 (16), 2393–2395.

41. Gerling, T.; Dietz, H., Reversible Covalent Stabilization of Stacking Contacts in DNA Assemblies. Angew Chem Int Ed Engl 2019, 58 (9), 2680–2684.

42. Willner, E. M.; Kamada, Y.; Suzuki, Y.; Emura, T.; Hidaka, K.; Dietz, H.; Sugiyama, H.; Endo, M., Single-Molecule Observation of the Photoregulated Conformational Dynamics of DNA Origami Nanoscissors. Angew Chem Int Ed Engl 2017, 56 (48), 15324–15328.

43. Martin, T. G.; Dietz, H., Magnesium-free self-assembly of multi-layer DNA objects. Nat Commun 2012, 3, 1103.

44. Valero, J.; Centola, M.; Ma, Y.; Skugor, M.; Yu, Z.; Haydell, M. W.; Keppner, D.; Famulok, M., Design, assembly, characterization, and operation of double-stranded interlocked DNA nanostructures. Nat Protoc 2019, 14 (10), 2818–2855.

45. Ouldridge, T. E.; Sulc, P.; Romano, F.; Doye, J. P. K.; Louis, A. A., DNA hybridization kinetics: zippering, internal displacement and sequence dependence. Nucleic Acids Research 2013, 41 (19), 8886–8895.

46. Snodin, B. E. K.; Romano, F.; Rovigatti, L.; Ouldridge, T. E.; Louis, A. A.; Doye, J. P. K., Direct Simulation of the Self-Assembly of a Small DNA Origami. Acs Nano 2016, 10 (2), 1724–1737.

47. Douglas, S. M.; Marblestone, A. H.; Teerapittayanon, S.; Vazquez, A.; Church, G. M.; Shih, W. M., Rapid prototyping of 3D DNA-origami shapes with caDNAno. Nucleic Acids Res. 2009, 37 (15), 5001–6.

48. Poppleton, E.; Bohlin, J.; Matthies, M.; Sharma, S.; Zhang, F.; Sulc, P., Design, optimization and analysis of large DNA and RNA nanostructures through interactive visualization, editing and molecular simulation. Nucleic Acids Research 2020, 48 (12).

49. Doye, J. P.; Fowler, H.; Prešern, D.; Bohlin, J.; Rovigatti, L.; Romano, F., … ; Snodin, B. E., The oxDNA coarse-grained model as a tool to simulate DNA origami. arXiv preprint 2020.

50. Skinner, G. M.; Kalafut, B. S.; Visscher, K., Downstream DNA tension regulates the stability of the T7 RNA polymerase initiation complex. Biophys J 2011, 100 (4), 1034–41.

