## Supplementary Information for "A rhythmically pulsing leaf-spring nanoengine that drives a passive follower"

### Materials and Methods

Methods

**Buffer systems.**

1× DNA buffer: 10 mM Tris·HCl, 50 mM NaCl, 10 mM $MgCl_2$ at pH 7.5.

1× origami buffer (NEOB): 50 mM Tris, 50 mM NaCl, 50 mM EDTA, 140 mM $MgCl_2$.

1× ligase buffer: 40 mM Tris·HCl, 10 mM $MgCl_2$, 10 mM dithiothreitol (DTT),

5 mM ATP at pH 7.8.

2× Precipitation Buffer (5% PEG 8000 (wt/vol) 5 mM Tris, 1 mM EDTA and 505 mM NaCl

5× transcription buffer (TB): 200 mM Tris·HCl (pH 7.9 at 25 °C), 50 mM DTT, 50 mM

NaCl and 10 mM spermidine.

**ODNs.** Oligodeoxynucleotides (ODN) staples used for the assembly of the DNA origami nano-engine structure were purchased from Eurofins Genomics (high-purity salt free). The M13mp18 (type 7249) scaffold used for the origami construction was purchased from Tilibit nanosystems GmbH. Chloroalkane-modified ODN (HaloTag Ligand O2 from the Biomers catalog) was purchased from Biomers as HPLC purified. ODNs with fluorophores (Cy3 and Cy5) or biotin as well as ODNs used for the transcribable DNA strand were obtained from Biomers and Microsynth as HPLC purified in dried state. LNA modified ODNs were purchased from Microsynth as HPLC purified ODNs. Upon arrival all ODNs have been diluted in MQ $H_2O$ to obtain a 100 µM concentrated solution by following manufacturer specifications.

**Assembly of the transcribable linker.** The transcribable DNA sequences were assembled from a subset of shorter 5’ phosphorylated ODNs (sequences can be found in the Supplementary data document #origami ODN#) that are designed to be partially complementary and overlapping so that they can be annealed in a temperature gradient and subsequentially ligated. The ODNs for the respective sequence are mixed in 5 µM concentration in 1 x ligase buffer with addition of NaCl in a final concentration of 20 mM. The samples were heated to 95 °C for 1 min to denature secondary structures and then annealed in a thermocycler with a temperature gradient from 60 °C to 15 °C during 75 min. After the annealing process Ligase (2 µl/150 µl, 10 U) was added and the sample was ligated over night at 15 °C. The Ligated product was separated from the not ligated ODNs and the ligase mixture utilizing Amicon Ultra 100 kDa size exclusion filter by spinning at 5000 rcf for 5 min. The buffer was exchanged to 1 x DNA buffer by subsequentially spinning for 5 more times by adding 500 µl off buffer each time. Samples were recovered by inverting the filter tubes into an empty reaction tube and spinning for 10 min at 3000 rcf. Concentration was determined by measuring the absorbance at 260 nm.

**Design NE DNA origami.** The origami was designed around the M13mp18 circular single stranded DNA scaffold of 7249 nucleotides in length utilizing the three-dimensional DNA designing tool Cadnano2 (https://cadnano.org). Staples have been optimized to have an average length and when possible optimized to follow design strategies presented in previous publications that described the $MgCl_2$ free assemblies of DNA origamis[43]. Three-dimensional representation of the design was obtained with the online Tool CanDo (https://cando-dna-origami.org/). The tool provides 3D structure predictions based on the cdano2 designs. The provided rmsf 3D maps have been used to optimize the structure to maximizes the overall origami stability. To ensure a high rigidity the stiff origami arm have been designed are formed by a 60 nm long 18 helixbundle in a honeycomb lattice arrangement. To permit movement in the DNA origami a compliant structural feature inspired by other works[20] has been inserted between the two stiff arms. The complaint region is made out of a 6 helix DNA “sheet” with a length of 84 nucleotides. The flexible double-stranded region is flanked by 6 single-stranded sequences that are part of the scaffold. To obtain the bend origami structure the 6 single-stranded sequences are designed to be 53 nucleotides long so that they are shorter than the complaint dsDNA sequence creating a tension in the hinge are that brings the origami to the desired angulated shape. From the optimized designs, atomic models are obtained through the CanDo tool and further used to confirm shape and dimension of the designed structure and arefurther used to refine the origami and introduce need modifications such as fluorophore, transcribable sequence, polymerase anchoring and biotin modifications. To introduce the chloroalkane modified staple, that serves as attachment point for the Halo T7 RNAP fusion protein, and the transcribable dsDNA we choose to select the more stable part of the origami that correspondsto the stiff origami arms at half of their length. In order to not reduce the overall stability of the origami, we have chosen to introduce modifications to the overall structure by choosing staples that already are nicked and are in the correct position where the modifications are needed. To ensure that the modification protrude normal to the surface of the origami, the neighboring staple is also extended out from the origami core and the design selected in such a way that the two protruding oligos form complementary duplex stem region. The double-stranded sequences that protrudes from the origami ensures that the sequence is orthogonal to the origami and extends away from the surface as long as the sequence is much shorter than the persistence length of DNA. Biotinylated ODNs have been introduced to the outside face of the origami in a tripod like conformation to increase the upright stability of the origami when anchored to surfaces via streptavidin (Sequences can be found in the Supplementary data document #origami ODN#).

**Assembly of the NE DNA origami.** The origami was assembled by combinig 13.3 nM of the M13mp18 scaffold with 10 EQ of each staple and 5 EQ of the transcribable dsDNA bridge in 1 x NEOB. The mixture is then divided into 50 µl aliquots into 500 µl reaction tubes and the origami annealed in a thermocycler. The aliquotation is performed to ensure that the samples are fully immerged into the thermoregulating element of the thermocycler Sequences and details for the different structures can be found in Supplementary Data Sets S1 and S2.

**Origami purification.** Structures are purified by precipitation of the origami structure in PEG containing buffer. The aliquoted origamis are combined in one 1.5 ml reaction tube and mixed 1:1 with the 2 x Precipitation buffer. The samples are spun at 16000 rcf for 30 min at 20 °C. The supernatant is removed with a pipette by making sure to not touch the pellet. Excess precipitation buffer is removed with thin strips of Whatman filter paper, always being careful not to touch the pellet. To the pellet 1 x OB is added. The pellet was suspended in 5 µl of 1 x NEOB for every 100 µl of assembly mixture. The samples were placed into an Eppendorf ThermoMixer C and shaken at 1500 rpm at 25 °C for at least 2h to fully resuspend the sample. Successful purification was confirmed by running 0.3 µl of resuspended sample on 1 % agarose gel.

**Expression and purification of Halo T7 RNAP fusion protein**: The Halo T7 RNAP fusion protein has been expressed and purified following the same protocol that has been utilized in another publication to express and purify the T7RNAP-Zif protein[22, 44] The differences in the procedure is the change of the expression plasmid in the Escherichia coli (strain BL21 DE3) with the plasmid pQE80HT-HaloTag-T7-RNAP (H= 6xHisTag, T=TEV site (tobacco etxh virus cleaving side), HaloTag=297 AA HaloTag protein tag, T7RNAP= 883 AA T7 RNA polymerase. In contrast to the already published protocol the TEV cleaving step is not performed. In brief the protein was expressed by starting from an overnight preculture at 37 °C in lysogeny broth medium with 50 µg/ml kanamycin. The preculture was diluted to an optical density at 600 nm (OD600) of 0.4 and when cells reached mid log phase of 0.6 (OD 600) expression was induced with addition of isopropyl β d thiogalactoside (IPTG) to a final con c of 0.5 mM and followed by incubation at 37 °C for 4h. Cells were collected by centrifugation at 4000 rpm. The cell pellet was resuspended in 15 ml lysis buffer (50 mM Tris, pH 7.8 at 4 °C, 300 mM NaCl, 10% glycerine, 20 mM imidazole and 1 mM $ZnCl_2$) and subsequentially the cells have been disrupted with a French press (1000 psi max, two rounds), spun for 20 min 20000 rpm at 4 °C and incubated with 1.5 ml of Ni-NTA agarose equilibrated bead matrix (Macherey Nagel) for 30 min and washed three times with the lysis buffer. The fusion protein was purification by affinity chromatography, then washed twice and eluted with elution buffer (lysis buffer containing 250 mM imidazole). The final purification step involved size exclusion chromatography (SEC) of the sample using a Superdex 200 column (GE Healthcare Life Sciences), fractions were collected, and the buffer was exchanged from dialysis buffer (Slide-A-Lyzer Dialysis Cassettes, 10K molecular weight cutoff) to storage buffer (50 mM Tris, pH 8.0, 0.1 mM EDTA, 300 mM NaCl and 50 % (vol/vol) glycerol).

**Transcription experiments.** The samples are prepared to obtain a final concentration of the Origami structure of 10 nM with addition of 2 EQ Halo T7 RNAP fusion. The Origami and the fusion protein are pre mixed in the right ratio in a premix with only addition of

TB but without diluting the sample to final concentration and incubated for 1 h in ice to favor the combination of the two components. To be able to follow the transcription the molecular beacon (MB) is added in a final concentration of 600 µM with addition of ribonuclease inhibitor ( 0.38 U/ul Recombinant RNasin® Promega). To start the transcription 2 mM of NTPs are added to the solution and the sample is diluted and brought to the final concentration in a mixture of 1 x NEOB and 1 x TB with addition of NaCl to a final additional concentration of 120 mM. The Fluorescence is monitored in a thermocycler over at least 3.5 h at 37 °C (Excitation wavelength 491 nm, Emission wavelength 517 nm). Sample volume loaded for each well is 20 µl for PerkineElmer ProxiPlate™-384 F Plus plates or 30 µl for Greiner Fluotrac 200 plates.

**MB signal calibration.** To estimate the amount of the transcript produced during the transcription experiments we calibrated the system using known amounts of an ODN (NLS 2) combined with the MB in the same buffer composition as for the transcription experiments. The sample is then treated as the transcription samples and the fluorescence is recorded over time. The obtained fluorescence signals are then used to generate a linear regression fit that is then used to estimate the amount of transcript generated during the transcription runs.

**Chloroalkane ODN competition experiments**. To test the competition due to the chloroalkane linker we tested preincubating the Halo-T7 RNApol equimolarly with the chloroalkane ODN for 1h on ice before adding it to the origami. To test if the chloroalkane ODN can displace the Halo-T7 RNApol from the origami we assemble the NE as described in the previous chapter and after the preincubation on ice of the NE we added 1, 2 or 5 EQ of the chloroalkane ODN to the origami solution and incubated for additionally 1h on ice before proceeding to prepare the samples for transcription as described in the previous chapter.

**Molecular dynamics simulations.** Multiple designs of the NE were simulated using the oxDNA coarse grain model for DNA origami. OxDNA is described in detail elsewhere[27-30], but briefly, it is an empirically-derived force field designed with DNA nanostructures in mind. The model has been shown to reproduce structural, kinetic and thermodynamic properties of DNA, including persistence length, strand displacement rates and free energy barriers between states, with reasonably high accuracy, while still being sufficiently coarse-grained to allow simulations of DNA origami at timescales of up to milliseconds[45-46]. Equilibrium simulations of the NE were carried out for 3 different designs: nNE, NE, and a version of nNE with two staples removed from the hinge (soft nicked NE, snNE). Starting configurations were obtained by exporting the CaDNAno[47] design file into oxDNA format using an in-house conversion script. Rigid body dynamics in oxView visualization tool[48] were used to bend the arms into a starting configuration. Relaxation was then performed using the method described in an ArXiv preprint[49]. After relaxation, the bridge was built using oxView's editing tools and a further round of relaxation performed. After the average energy per particle stabilized around -1.5 su, the structures were equilibrated with production conditions for a further 2.5e8 simulation steps to allow equilibration of the angle distribution (relaxation step number determined by initial exploratory simulations). Equilibrium simulations were carried out for 1e9 oxDNA simulation steps with a timestep of integration of 0.003, a temperature of either 23° C or 37° C was imposed using an Andersen-like thermostat. Configuration snapshots were saved for analysis every 5e5 steps giving 2000 configurations per simulation which were verified to be well decorrelated. To identify the effects of secondary structure in the flexure region, a second set of equilibrium simulations was performed with the base type of the nucleotides in the single-stranded region of the flexure set to non-interacting (no structure, NS). These simulations were carried out with the same parameters as the simulations where secondary structure was allowed to form.

Closing rates were measured by running either 3 (snNE) or 10 (nNE, nNE-NS, NE and NE-NS) replicates of simulations with external forces added. The simulations were started from the final configuration of one replicate of the associated equilibrium simulation and run for 1e7 steps with snapshots saved every 1e3 steps for a total of 2000 configurations per simulation. A constant force of 16 pN (based on the typical value of tension exerted by a polymerase on a duplex DNA[50]) was applied between the nucleotide where the RNA polymerase is covalently linked to the first nucleotide in the first stop sequence in the bridge with a cutoff radius of 10.35 nm (the radius of T4 polymerase)[31]. Other parameters were the same as in the equilibrium simulations. One additional simulation was performed for each design where the force was applied for 1e9 steps to allow the structure to equilibrate in the closed position.

Opening rates were measured by running either 3 (snNE) or 10 (nNE, nNE-NS, NE and NE-NS) replicates starting from the final configuration of the equilibrated pulling simulation. Each simulation was run for 2e7 steps with snapshots saved every 1e3 steps for a total of 4000 configurations per simulation. In addition to the full NE, simulations were also performed where the bridge was deleted (NE-NB and NE-NS-NB), allowing the structure to open under only the influence of the flexure. Other parameters were the same as in the equilibrium simulations.

Simulations were aligned and mean structures obtained from the equilibrium simulations using oxdna_analysis_tools[48] and movies of the trajectories produced using oxView[48]. To get the spring constant and opening/closing rates, a linear regression was performed on the point clouds corresponding to the arms of each hinge and the angle between the arms calculated for each snapshot. The spring constant was then estimated using the equipartition theorem for a simple harmonic oscillator

$$k(x-x_0)^2 = ½ K_bT$$

Where k is the spring constant, x and $x_0$ are the displacement and the average position, and $K_bT$ is the thermal energy. These values were converted from oxDNA simulation units to pN/deg using the conversion factors available in the oxDNA model introduction.

Opening and closing rates were calculated using a linear regression to the angle traces over the simulations. As with spring constant, these rates were converted to deg/ns using the conversion factors. It should be noted, however that due to coarse-graining and the increased diffusion constant used by the oxDNA model, exact experimental time correlation with processes in coarse grained is often under-estimated by direct unit conversion. Past research on DNA hybridization suggests that rates calculated via direct unit conversion may be $10^4$-fold lower than actual time correspondence[46]. Significance of distributions was determined using a two-tailed Kolmogorov-Smirnov test.

**AFM imaging:** atomic force microscopy is performed on a JPK NANOWIZARD 3. Scanning was performed with ACTA-50 SPM probes, made out of Si (N-type) with 0.01-0.025 ohm/cm. The the used cantilevers are 125 µm long, 30 µm wide and 4 µm tall with a f: 200-400 kHz resonance frequency and a spring constant k: 13-77 N/m and an AL coating on the reflex side. Samples are prepared by diluting the samples to 10 nM concertation in 1 x Origami Buffer. Two microliters of the samples are placed on freshly cleaved mica inside circle (3 mm in diameter) drawn with a thin tip marker pen to spatially confine the droplet and to easily find the deposition area afterwards. The sample is incubated on the surface for 2 min and then washed three times by slowly dripping 200 µl on the mica surface. The surface is dried with a gentle airstream and the samples imaged in intermitting contact mode on 1 – 2 µm squares with a 512 x 512-pixel resolution.

**Transmission Electron Microscopy (TEM)**. The samples was applied at a concentration of 1 to 10 in a volume of 3 µl on a 5 nm continuous carbon (Leica EM ACE600) coated TEM grids (Cu 3 mm 400 MESH TEM GRID, SPI-GIDS™). The sample was stained using a 2% uranyl formate solution. Images were taken in Low-dose mode using a FEI Tecnai Spirit Bio-Twin Microscope at 120kV equipped with a Gatan US4000 4k x 4k CCD camera. The magnification used was 30kx (pixel size of 3.80Å) for angle distributions and 68kx (pixel size 1.66) for details.

**Angle Measurement from TEM images.** To determine the angle distribution several TEM images have been scanned using the open-source imaging software Fiji (imagej.net/software/fiji/), a package distribution of ImageJ2. The Angle Tool was used to measure the angle that is formed between the two stiff arms of one origami structure by aligning the line from the Angle Tool to the helixes that are visible due to slight differences in the electron density of the origami matrix. All structure with a clearly identifiable, correct and complete structure have been measured, without making any distinction on the visible angle. Structures that are broken, showed signs of missing parts or were not clearly lying flat on the surface were ignored in the angle measurements assuring so that only intact origamis have been measured. In case of the "Driver"-"Follower" only construct that clearly show the connected origami structures are considered for the angle measurement. The Angle of the D and the F units are measured for each structure.

**Driver and Follower constructs**. Driver and Follower origami are assembled and purified separately as described in the origami assembly section with the only exception that the connecting ODN are added in twice the amount compared to the other ODNs. The structures are purified as described in the origami section with precipitation in PEG Precipitation Buffer. The purified structures are equimolarly combined without further diluting the system and incubated at 30 °C over 2 h to favor the combination of the two structures. After the first incubation the samples are subsequentially diluted for the further application.

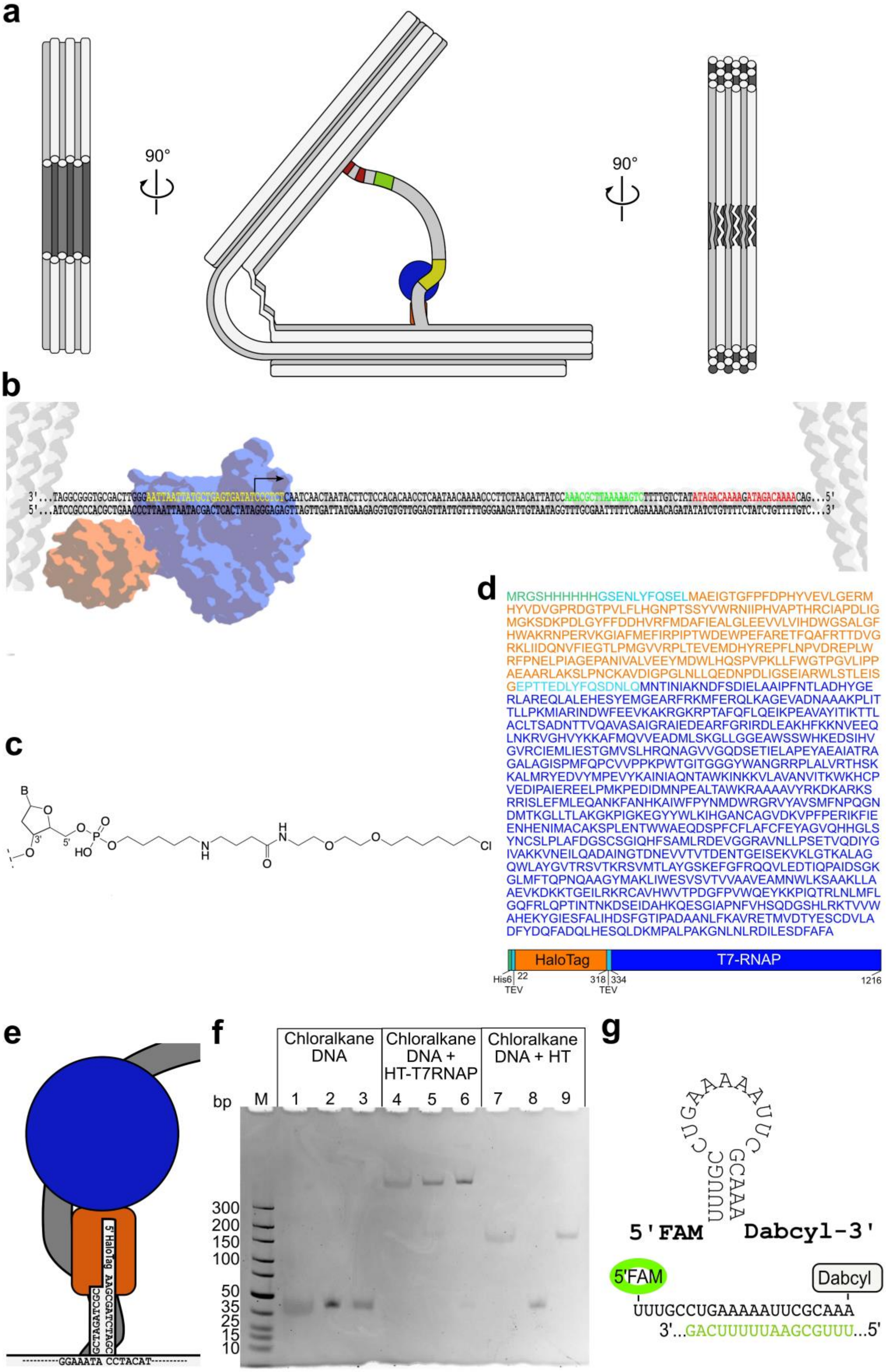
a
90°
90°
b
c
B
3'
5'
O
P
HO
H
N
O
Cl
d
MRGSHHHHHHGSENLYFQSELMAEIGTGFPFDPHYVEVLGERM
HYVDVGPRDGTPVLFLHGNPTSSYVWRNIIPHVAPTHRCIAPDLIG
MGKSDKPDLGYFFDDHVRFMDAFIEALGLEEVVLVIHDWGSALGF
HWAKRNPERVKGIAFMEFIRPIPTWDEWPEFARETFQAFRTTDVG
RKLIIDQNVFIEGTLPMGVVRPLTEVEMDHYREPFLNPVDREPLW
RFPNELPIAGEPANIVALVEEYMDWLHQSPVPKLLFWGTPGVLIPP
AEAARLAKSLPNCKAVDIGPGLNLLQEDNPDLIGSEIARWLSTLEIS
GEPTTEDLYFQSDNLQMNTINIAKNDFSDIELAAIPFNTLADHYGE
RLAREQLALEHESYEMGEARFRKMFERQLKAGEVADNAAAKPLIT
TLLPKMIARINDWFEEVKAKRGKRPTAFQFLQEIKPEAVAYITIKTTL
ACLTSADNTTVQAVASAIGRAIEDEARFGRIRDLEAKHFKKNVEEQ
LNKRVGHVYKKAFMQVVEADMLSKGLLGGEAWSSWHKEDSIHV
GVRCIEMLIESTGMVSLHRQNAGVVGQDSETIELAPEYAEAIATRA
GALAGISPMFQPCVVPPKPWTGITGGGYWANGRRPLALVRTHSK
KALMRYEDVYMPEVYKAINIAQNTAWKINKKVLAVANVITKWKHCP
VEDIPAIEREELPMKPEDIDMNPEALTAWKRAAAAVYRKDKARKS
RRISLEFMLEQANKFANHKAIWFPYNMDWRGRVYAVSMFNPQGN
DMTKGLLTLAKGKPIGKEGYYWLKIHGANCAGVDKVPFPERIKFIE
ENHENIMACAKSPLENTWWAEQDSPFCFLAFCFEYAGVQHHGLS
YNCSLPLAFDGSCSGIQHFSAMLRDEVGGRAVNLLPSETVQDIYG
IVAKKVNEILQADAINGTDNEVVTVTDENTGEISEKVKLGTKALAG
QWLAYGVTRSVTKRSVMTLAYGSKEFGFRQQVLEDTIQPAIDSGK
GLMFTQPNQAAGYMAKLIWESVSVTVVAAVEAMNWLKSAAKLLA
AEVKDKKTGEILRKRCAVHWVTPDGFPVWQEYKKPIQTRLNLMFL
GQFRLQPTINTNKDSEIDAHKQESGIAPNFVHSQDGSHLRKTVVW
AHEKYGIESFALIHDSFGTIPADAANLFKAVRETMVDTYESCDVLA
DFYDQFADQLHESQLDKMPALPAKGNLNLRDILESDFAFA
HaloTag
T7-RNAP
His6
22
TEV
318
334
TEV
1216
e
5'HaloTag AAGCGATCTAGC
GCTAGATCGC
GGAAATA CCTACAT
f
Chloralkane DNA
Chloralkane DNA + HT-T7RNAP
Chloralkane DNA + HT
bp
M
1
2
3
4
5
6
7
8
9
300
200
150
100
50
35
25
15
10
g
5'FAM
Dabcyl-3'
5'FAM
Dabcyl
UUUGCCUGAAAAAUUCGCAAA
3'...GACUUUUUAAGCGUUU...5'

**Suppl. Figure S1**. Additional design features of the leaf-spring NE (**a**) Cross section of the NE with the leaf-spring (left panel), a side-view (middle) that shows the ssDNA template and the location of the T7 promotor (yellow), the sequence coding for a binding site for a molecular beacon to determine transcription yields (green), and the two terminator sequences (red). The right panel shows the location of the six ssDNA sequences, (**b**) Sequence of the dsDNA transcription template. Yellow: T7 promotor, green: sequence coding for molecular beacon-binding (green), red: terminator sequence, (**c**) Chemical structure of the halogenated 5'-end of the protruding HT-T7RNAP attachment staple, (**d**) Primary amino acid sequence (upper panel) and design (lower panel) of the HT-T7RNAP fusion protein, (**e**) Design and sequence of the protruding staple containing the 5'-halogenated attachment site for HT-T7RNAP, (**f**) Chloroalkane DNA connection to the HaloTag (HT) enzyme. Lanes 1-3: Chloralkane DNA in (1) origami buffer, (2) $H_2O$, (3) Origami buffer + EDTA; lanes 4-6: Chloralkane DNA + HT-T7RNAP in (4) origami buffer, (5) $H_2O$, (6) Origami buffer + EDTA; lanes 7-9: Chloralkane DNA + HT in (7) origami buffer, (8) $H_2O$, (9) Origami buffer + EDTA. The Halo enzyme alone in absence of $MgCl_2$ cannot bind to the chloroalkane modified DNA. In presence of the buffer the protein can bind to the DNA and the connection is covalent and strong enough that even after addition of EDTA to remove the $MgCl_2$ the connection of DNA and protein is maintained. In the fusion protein of Halo T7 RNA pol this phenomenon is not present, very likely due to the affinity of the polymerase towards DNA that probably increases the affinity of the fused Halo tag towards the DNA, (**g**) Sequence, secondary structure, and labels (5'-FAM, 3'-Dabcyl) of the molecular beacon RNA that detects the green sequence in the RNA generated during transcription.

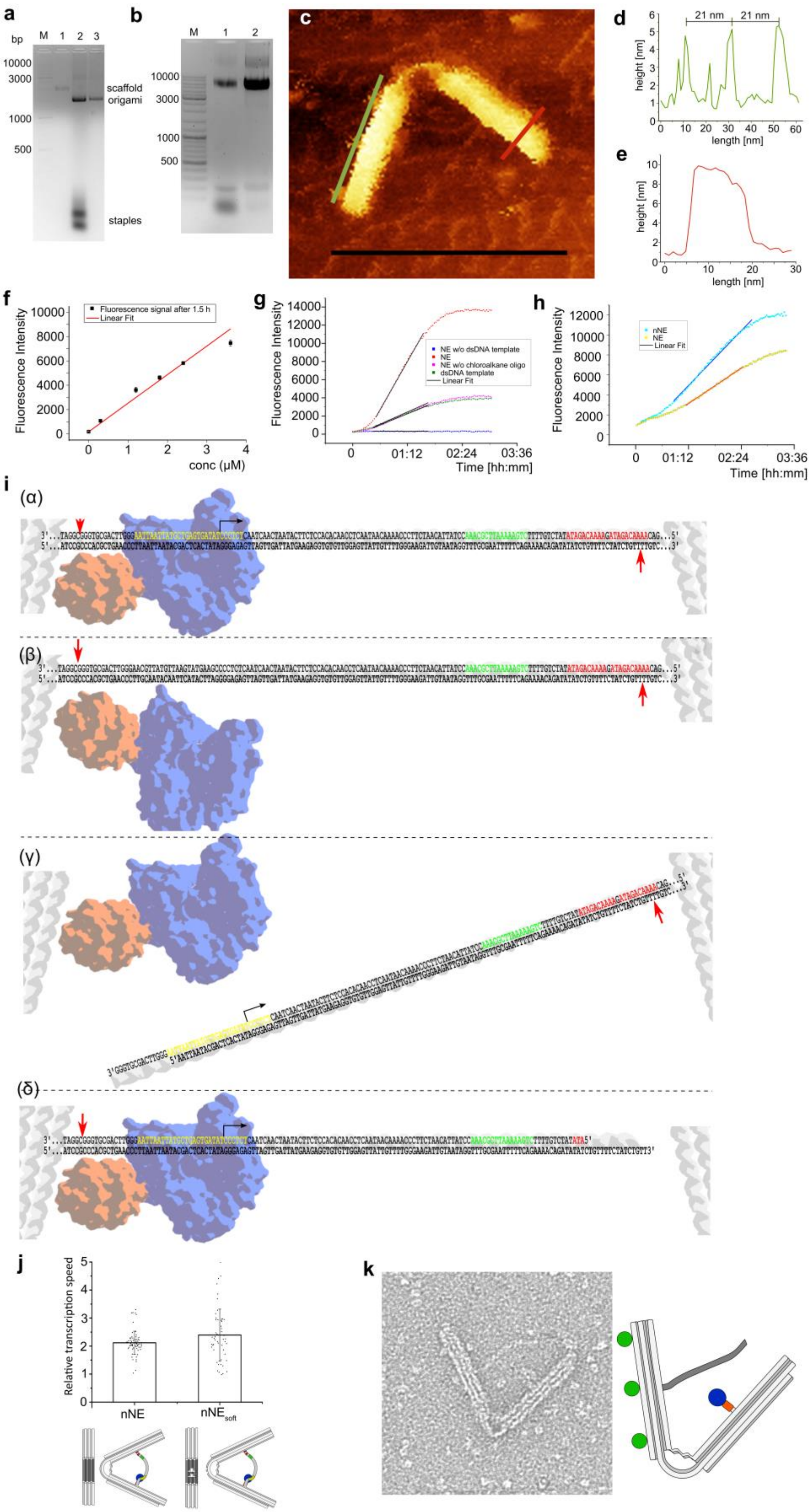
a
bp
M 1 2 3
10000
3000
1000
500
scaffold
origami
staples
b
M 1 2
10000
3000
1000
500
c
d
21 nm
21 nm
height [nm]
length [nm]
e
height [nm]
length [nm]
f
Fluorescence signal after 1.5 h
Linear Fit
Fluorescence Intensity
conc (µM)
g
NE w/o dsDNA template
NE
NE w/o chloroalkane oligo
dsDNA template
Linear Fit
Fluorescence Intensity
Time [hh:mm]
h
nNE
NE
Linear Fit
Fluorescence Intensity
Time [hh:mm]
i
(α)
(β)
(γ)
(δ)
j
Relative transcription speed
nNE
nNE soft
k

**Suppl. Figure S2**. Assemby of the NE and AFM analysis of the geometry of the NE. (**a**) Filter purification of the origami. Lane 1: M13MP18 scaffold, lane 2: assembled origami unpurified, lane 3: 100 kDa filter-purified origami, (**b**) PEG-purification of origami. Lane 1: Assembled unpurified origami, lane 2: PEG-purified origami, (**c**) detailed AFM image of the origami structure. The green line spans the streptavidin molecules attached to the respective origami-arm and marks the height measurement shown in (**d**). The red line marks the cross-section of the opposing origami-arm and marks the height measurement shown in (**e**), (**d**) height profile of the green line shown in (**c**) confirms the distance of the streptavidin molecules on the respective origami-arm to be spaced exactly 21 nm as designed, (**e**) height profile of the red line shown in (**c**) to determine the cross-section of the opposing origami-arm to be exactly 9 nm as designed, (**f**) calibration curve of the fluorescence intensity (F.I.) of the molecular beacon (MB) as a function of the concentration of the added complementary oligonucleotide. The measuring time was 1.5 h because the F.I. stabilized after that time without significant further photobleaching, (**g**) exemplary transcription curves of constructs 1a (blue), 2a (green), 3a (magenta), and 4a (red) shown in **Fig. 2a** with linear fit of the linear parts of the curves, (**h**) exemplary transcription curves of constructs 1b (yellow) and 2b (cyan) shown in **Fig. 2b** with linear fit of the linear parts of the curves, (**i**) Detailed representation of the various forms of template dsDNA used in this study; (α) red arrows: position of the two nicks in the template dsDNA, (β) red arrows: the same without the promotor region, (γ) dsDNA, attached only on the opposite side of the HT-T7RNAP; red arrow: nick-position, (δ) dsDNA, attached only next to the HT-T7RNAP; red arrow: nick position. Importantly, to avoid that the single stranded nicks weakens the stability of the template DNA and cause detachment of the DNA from the origami by forces generated during the pulling by the polymerase and/or during the formation of the transcription bubble, the template strand has no single nicks upstream of the promoter region while the nick is placed in the coding strand. Downstream of the promoter region the single stranded nick has been placed into the template strand while the coding strand is fully attached into the origami. This design avoids that the two ss nicks in the coding strand lead to loss of the DNA-DNA duplex by formation of a DNA-RNA as the amount of RNA increases over time. Both, leading and coding strands, are anchored to the origami; their hybridization is favored even when the RNA transcript levels increase, (**j**) Relative transcription speed of nNE (left) and $nNE_{soft}$ (right). Error bars: S.D., n ≥ 52. (**k**) representative TEM image (upper panel) of construct 1d shown in **Fig. 2d** indicates electrostatic repulsion of the dsDNA template strand in case where the strand is not anchored at its anchoring point next to the HT-T7RNAP, which hampers efficient transcription. Lower panel: cross section of this construct.

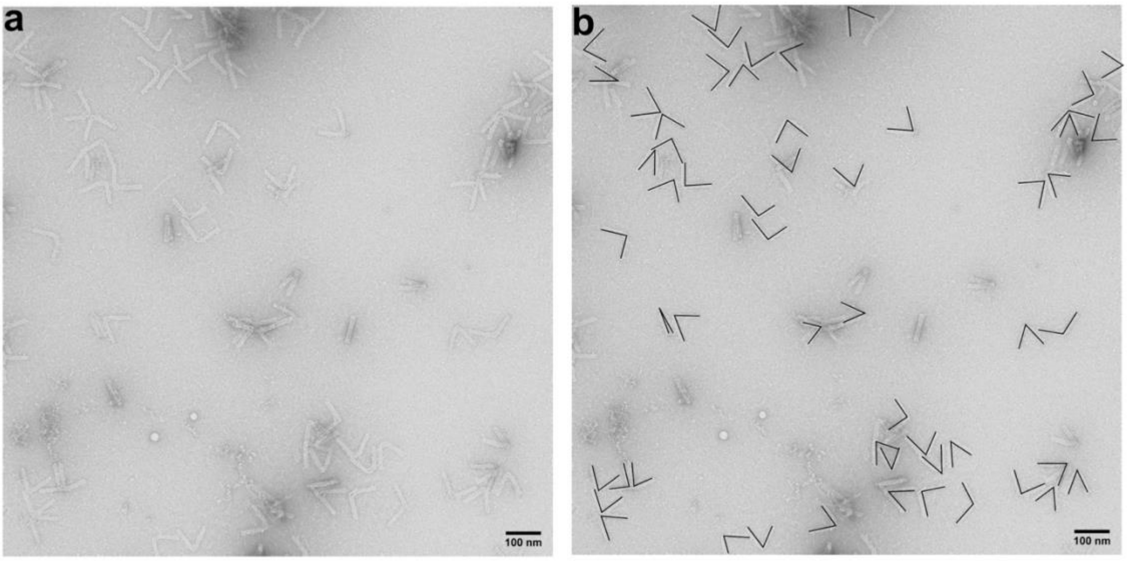


**Suppl. Figure S3**. Exemplary angle measurement under transcription conditions. (**a**) Example of raw data on a TEM image section. For angele measurements, only structures that lied flat on the surface were considered. These were identified based on visibility and correct form of the hinge structures as well as clear visibility of the axial dsDNA patterns in the origami arms. (**b**) Angle measurement on the same section. Angles were measured by using the software imageJ with the in FIJI processing package and the “angle”-tool. For reproducibility the straight lines corresponding to the DNA helices in the origami-arms were used to guide the proper alignment of the angle-tool. The central point of the angle tool was then adjusted until all the lines of the angle tool and the dsDNA in the structure matched (lower panel). Structures that had no clearly visible compliant hinge region or showed clear damages in the origami structure were excluded. Structures without visible damage that showed extremely narrow or wide angles were included in the measurement. Scale bar: 100 nm.

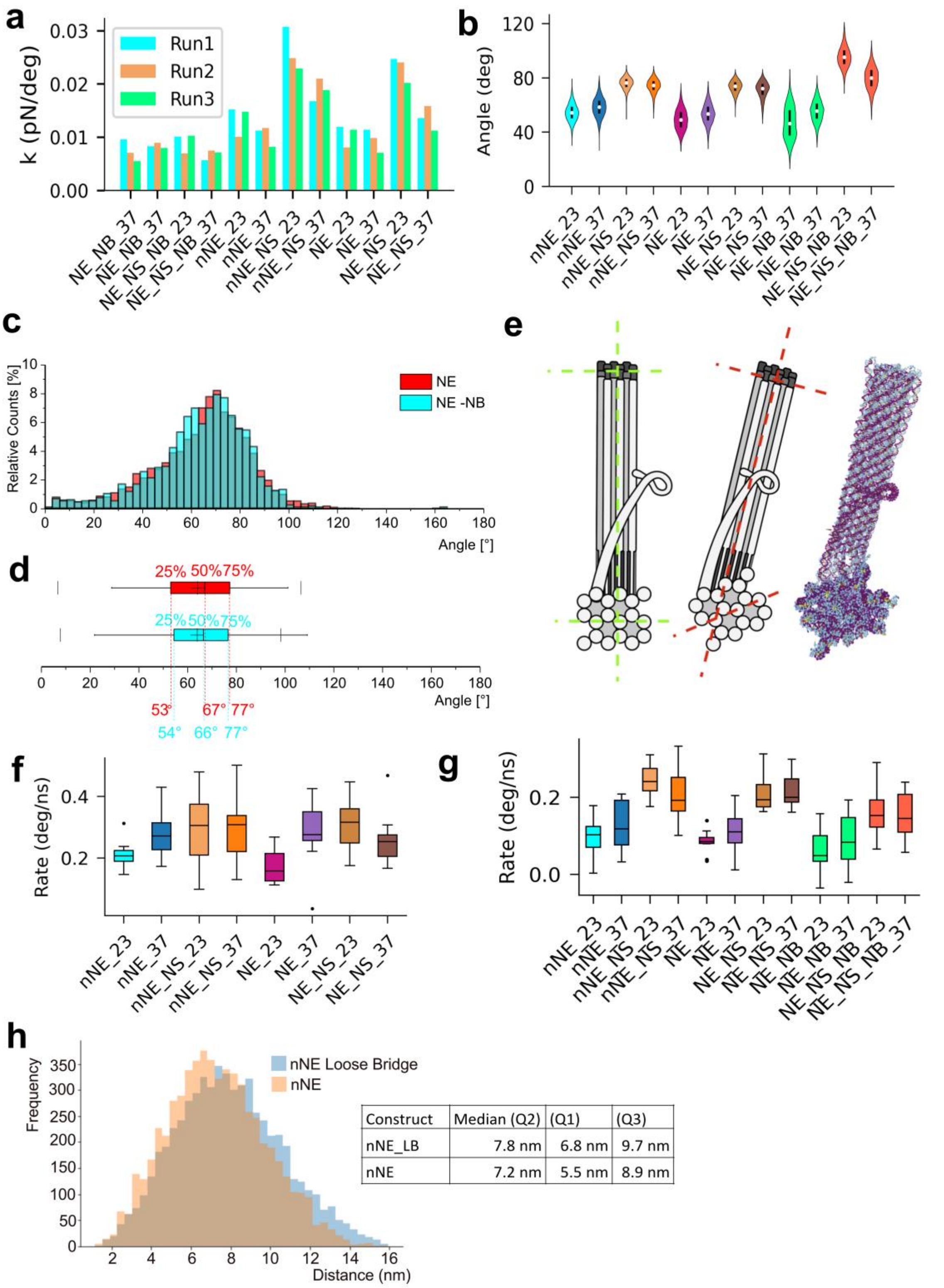


| Construct | Median (Q2) | (Q1) | (Q3) |
|---|---|---|---|
| nNE_LB | 7.8 nm | 6.8 nm | 9.7 nm |
| nNE | 7.2 nm | 5.5 nm | 8.9 nm |

**Suppl. Figure S4.** Comparison of hinge properties of simulations run at 23° and 37° C. **(a)** Calculated spring constant of each design at 23° and 37° C. Only NS structures with an intact dsDNA bridge show a significant difference between the two temperatures, demonstrating the influence of stable secondary structures on angle distribution, **(b)** Angle-distribution in triplicate equilibrium simulations. For most structures, the angle distribution did not change much when the temperature was increased. A slight increase in average angle for simulations with secondary structures in the flexure and slight decrease in average angle for simulations where secondary structure was inhibited was observed. This is consistent with the hypothesis that secondary structures limit the opening angle of the structures with secondary structure while the extended single strands behave more like entropic springs which become more flexible as temperature increases, **(c, d**) To confirm that the angle distribution of the structure depends primarily on the features of the hinge and not on other features as for example the transcribable dsDNA strands we confronted the distribution of angles measured from TEM images of NE (red, n=5135) with the angle distribution of the NE-NB missing the dsDNA template strand (cyan, n=1382), (**c**) The two angle distributions are highly comparable (p = 0.6). The box plot in (**d**) with whiskers of size 1.5 times with first, second (median) and third quartile indicated with dashed lines (red for NE and cyan for NE-NB). Thin cross: average values 64° ±20° for NE and 64° ±19° for NE-NB, (**e**) Relaxation of the NE origami structure simulation indicate that the origami arms bent out of plane. Left panel: designed structure, middle panel: relaxed simulation, right panel: direct view on one of the stiff 18 HB stiff origami arms while the second arm extends towards the top of the structure. The origami arms clearly show a longitudinal twist in the top view of the 18 HB and are beni out of the vertical plane of the arm that extends towards the top. The reference lines are placed in green in the designed structure and in red in the simulated structure to illustrate the amount of bending. Since the dsDNA helixes in the origami arms are used as references, a distortion of the surface-deposited structure results in slight systematic difference compared to how the angles are measured in the simulated structure. This difference likely explains the systematic difference in the angle distributions. **(f)** Calculated pulling rates under 16 pN applied force between the polymerase attachment point and the terminator sequence on the dsDNA bridge for each design at 23° and 37° C. The only significant difference between the two temperatures was observed for nNE and NE, where decreasing the number of base pairs in the flexure significantly increased the rate at which the leaf-spring was able to close, **(g)** Calculated re-opening rates after being closed under 16 pN force. This is a process which relies mostly on brownian motion to return to the relaxed state of the NE. Unsurprisingly, we see an increase in average rate and an increase in variance for the structures where secondary structures can form in the flexure and a decrease in average rate and no trend in variance for structures where no structure was permitted in the flexure, (**h**) Frequency of time that the promoter region spends at different distances to the point where the polymerase is anchored to the origami. The radius representing the attached polymerase has been estimated to be 7.5 nm to have an encounter of promoter region and polymerase. The frequency at which the nNE construct (orange curve, nNE) spends within this ideal distance bubble is higher then the case where the transcribable dsDNA strand is attached only next to the polymerase (blue curve, nNE_LB ). The median value for the nNE is 7.2 nm with the Q1 = 5.5 nm and Q3 = 8.9 nm. In case of the attachment only next to the polymerase the median is bigger than the 7.5 nm reference radius at median value of 7.8 nm with Q1 = 6.8 nm and Q2 = 9.7 nm.

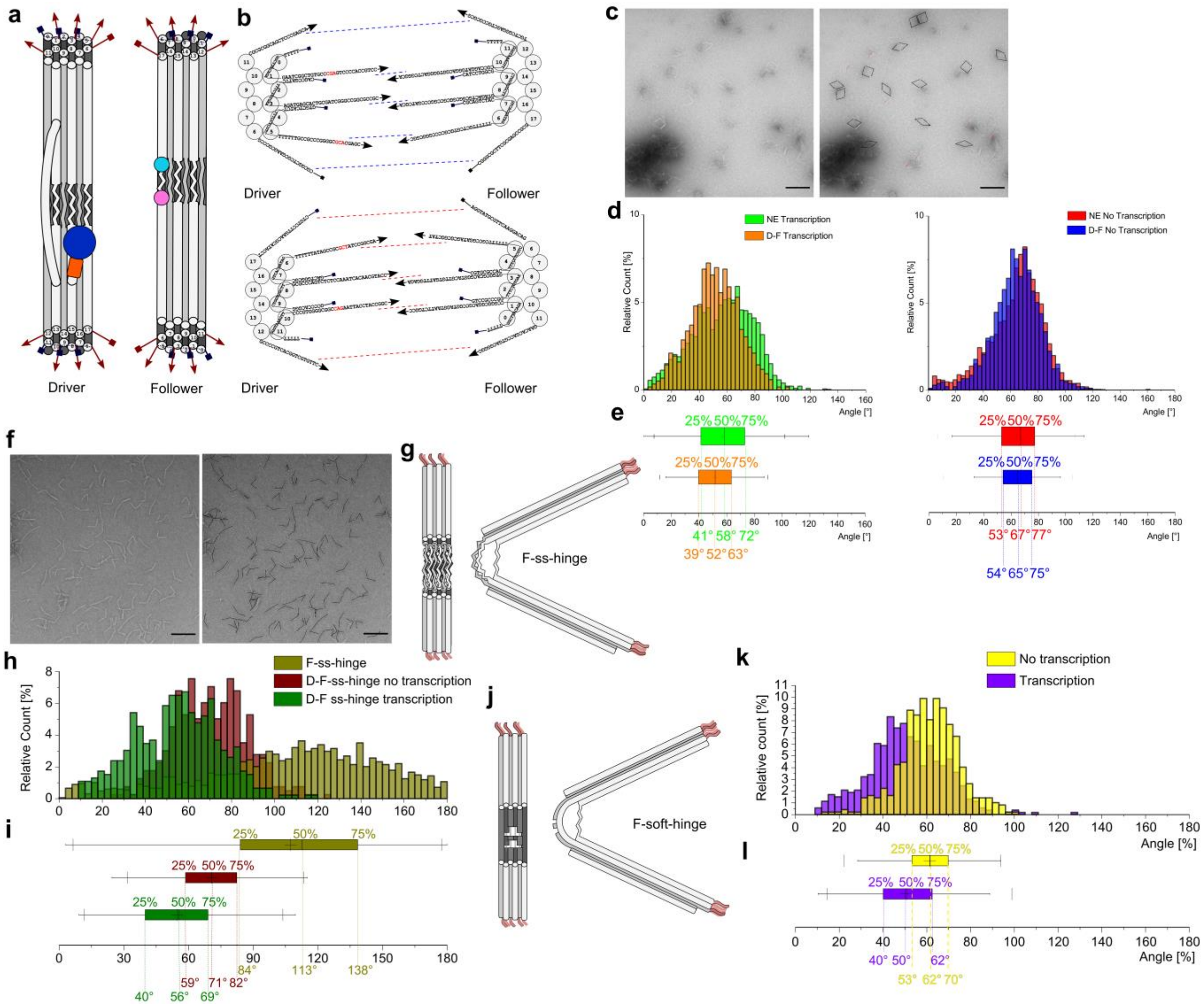
a
Driver
Follower
b
Driver
Follower
Driver
Follower
c
d
NE Transcription
D-F Transcription
NE No Transcription
D-F No Transcription
Relative Count [%]
Angle [°]
e
25% 50% 75%
41° 58° 72°
39° 52° 63°
53° 67° 77°
54° 65° 75°
f
g
F-ss-hinge
h
F-ss-hinge
D-F-ss-hinge no transcription
D-F ss-hinge transcription
Relative Count [%]
i
25% 50% 75%
84° 113° 138°
59° 71° 82°
40° 56° 69°
j
F-soft-hinge
k
No transcription
Transcription
Relative count [%]
Angle [%]
l
25% 50% 75%
40° 50° 62°
53° 62° 70°

**Suppl. Figure S5.** Schematic front view of the Driver and Follower units, (**a**) Helices are numbered from 0 to 17 as a guide to better follow the positioning of the single stranded overhangs. The numbering of the helices is equivalent in both origami structures, please note how it is necessary to flip one of the two units upside down to be able to join the structures, (**b**) detailed DNA sequences of the overhangs for the Driver (left) and for the Follower (right). Bases labeled in red in the Driver overhang sequences indicate LNA. TEM images have been used to measure the angle distribution of the different constructs, (**c**) shows an example of TEM image used to determine the angle of the D-F complex on the left and with the overlayed black line to indicate how the angle were measured. Not properly joined structures or single origamis have been ignored during the angle measurements (red strikethrough). When measuring the angle distribution from TEM images (**d**) and comparing the distribution of NE (red, n = 5135) with the angle distributions of only complete D-F (blue, n = 1074) in absence of transcription it is noticeable how the average angle remains unchanged, being 64° ± 20° for NE and 64° ± 17° (Error: S.D.) but the curve is less skewed and more symmetric in case of the D-F complex. The same effect is appreciable also during transcription (NE in green, n = 3266; D-F in orange, n = 1190). In the NE sample the average angle distribution drops to 57° ± 22° that is comparable to the average value of the D-F complex of 57° ± 17° but the curve is narrower in the second case. The boxplots of the distribution (**e**), confirm that in case of the D-F complex the angle distribution is narrower and less skewed. The stabilizing effect on the angle distribution is particularly appreciable in the case of the D-F-no-hinge (**e**). The example TEM image for the F-no-hinge structure (**f**) nicely shows how large the range of angles is when the double stranded structure is missing in the flexure region making it impossible to obtain the correct structure that determines the angulated form of the origami structure. The F-no-hinge (**g**) constructs shows a verry wide and flat distribution (**h**) (dark yellow, n = 1682) with obtuse angle of 107° ± 41°. When combining the F-no-hinge origami with the D unit the angle is reduced to 71° ± 17° with a narrower distribution (wine, n = 398). In case of transcription the average distribution angle shifts to 55° ± 20° getting slightly larger as expected for the transcription sample (olive, n = 462). The Boxplots (**i**) show clearly how the distribution for the F-no-hinge (dark yellow) is widely spread over a great range of angles, having a difference between Q3 and Q1 of 54° while in the D-F this range is reduced to 23° in absence of transcription (wine) and 29° in presence of transcription (olive). The change in angle distribution during transcription in the D-F structure is also appreciable in the presence of F-soft-hinge (**j**) in the complex (**k**). The

D-F-soft-hinge complex shows an average angle of 61° ± 14° (yellow, n = 496) in the no transcription case and 51° ± 17° in the transcription sample (violet, n = 528), (**I**) the boxplot graphically shows how the distribution clearly shifts towards more acute angles in the case of transcription compared to the no transcription sample. Errors are S.D. and the boxplots show Q1, Q2, Q3 in the box with whiskers 1.5 times the box size. Thin crosses in the boxplots indicate the average angle while the thin vertical lines indicate the 1% and 99% percentile of the distribution.

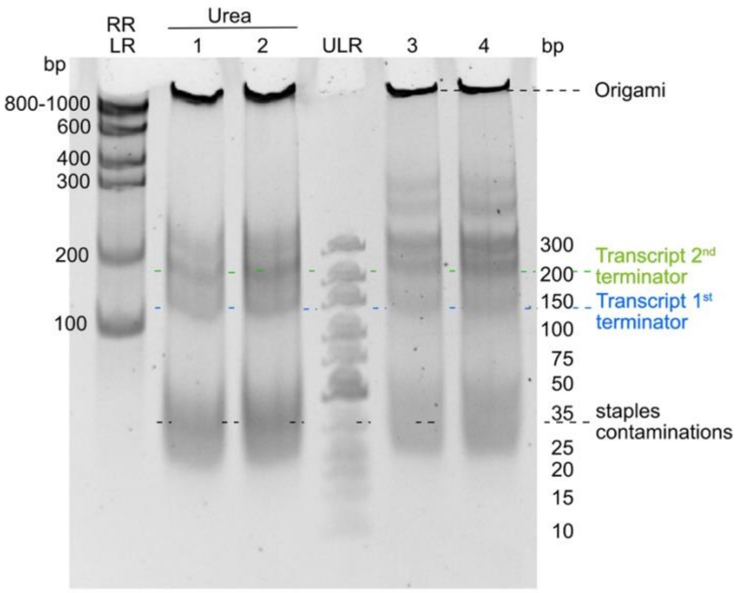


**Suppl. Figure S6**. Amounts of RNA observed by polyacrylamide gel electrophoresis (PAGE) for the D-F-pair. Lane 1: D-F batch 1 + urea, lane 2 D-F-batch 2 + urea, lane 3: D-F batch 1, lane 4: D-F batch 2. RR-LR: RiboRuler Low Range ss RNA ladder; ULR: Ultra low range dsDNA ladder.
